# Mathematical modeling of plant cell fate transitions controlled by hormonal signals

**DOI:** 10.1101/830190

**Authors:** Filip Z. Klawe, Thomas Stiehl, Peter Bastian, Christophe Gaillochet, Jan U. Lohmann, Anna Marciniak-Czochra

**Affiliations:** Institute of Applied Mathematics, Heidelberg University, Heidelberg, Germany; Interdisciplinary Center for Scientific Computing, Heidelberg University, Heidelberg, Germany; Bioquant Center, Heidelberg University, Heidelberg, Germany; VIB-UGent Center for Plant Systems Biology, Ghent University, Ghent, Belgium; Department of Stem Cell Biology, Centre for Organismal Studies, Heidelberg University, Heidelberg, Germany

**Keywords:** Arabidopsis thaliana, shoot apical meristem, mathematical model, stem cells, reaction-diffusion equations, growing domain, hormonal feedback, WUSCHEL, CLAVATA, HECATE

## Abstract

Coordination of fate transition and cell division is crucial to maintain the plant architecture and to achieve efficient production of plant organs. In this paper, we analysed the stem cell dynamics at the shoot apical meristem (SAM) that is one of the plant stem cells locations. We designed a mathematical model to elucidate the impact of hormonal signaling on the fate transition rates between different zones corresponding to slowly dividing stem cells and fast dividing transit amplifying cells. The model is based on a simplified two-dimensional disc geometry of the SAM and accounts for a continuous displacement towards the periphery of cells produced in the central zone. Coupling growth and hormonal signaling results in a non-linear system of reaction-diffusion equations on a growing domain with the growth velocity depending on the model components. The model is tested by simulating perturbations in the level of key transcription factors that maintain SAM homeostasis. The model provides new insights on how the transcription factor HECATE is integrated in the regulatory network that governs stem cell differentiation.

**Summary:** Plants continuously generate new organs such as leaves, roots and flowers. This process is driven by stem cells which are located in specialized regions, so-called meristems. Dividing stem cells give rise to offspring that, during a process referred to as cell fate transition, become more specialized and give rise to organs. Plant architecture and crop yield crucially depend on the regulation of meristem dynamics. To better understand this regulation, we develop a computational model of the shoot meristem. The model describes the meristem as a two-dimensional disk that can grow and shrink over time, depending on the concentrations of the signalling factors in its interior. This allows studying how the non-linear interaction of multiple transcription factors is linked to cell division and fate-transition. We test the model by simulating perturbations of meristem signals and comparing them to experimental data. The model allows simulating different hypotheses about signal effects. Based on the model we study the specific role of the transcription factor HECATE and provide new insights in its action on cell dynamics and in its interrelation with other known transcription factors in the meristem.

## Introduction

Tissue function is an effect of the cooperation of multiple specialized cell types. To establish, maintain and regenerate tissues, cell production and fate specification have to be orchestrated in a robust and well-defined manner. Perturbations of the underlying control mechanisms may reduce the ability of the organism to adapt to changing environmental conditions.

Plants continuously generate new organs such as leaves, roots and flowers. For this purpose they maintain pools of stem cells which remain active during the whole life of the plant. The plant stem cells are located in specialized tissues, referred to as *meristems*. The accessibility of meristems to live-imaging and the repetitive formation of identical organs, such as leaves, make plants an attractive system to study the regulatory cues underlying cell production and fate transition.

Stem cell proliferation and fate choice have a direct impact on the architecture of the plant and its reproductive fitness. These vital functions require meristems to be robust with respect to perturbations such as injuries or environmental fluctuations. In case of agricultural plants, meristem dynamics are linked to the crop yield, therefore, a better understanding meristem regulations is of practical importance [1, 2].

In this paper we focus on the stem cell dynamics in the shoot apical meristem (SAM) that is responsible for formation of all above ground structures. The SAM is a curved structure and its geometry can be approximated either by a spherical cap [3] or by a paraboloid [4]. Stem cells are located in the central zone (CZ) surrounded by transit amplifying cells in the peripheral zone (PZ). Newly formed but still immature organs, so-called primordia, separate from the meristem at the outer boundary of the PZ. A specialized cell population, the so called organizing center (OC), is located below the CZ and produces signals required for maintenance of the stem cell fate, see [5]. The location of the SAM and its morphology is summarized in Fig. 1.

**Figure 1:**
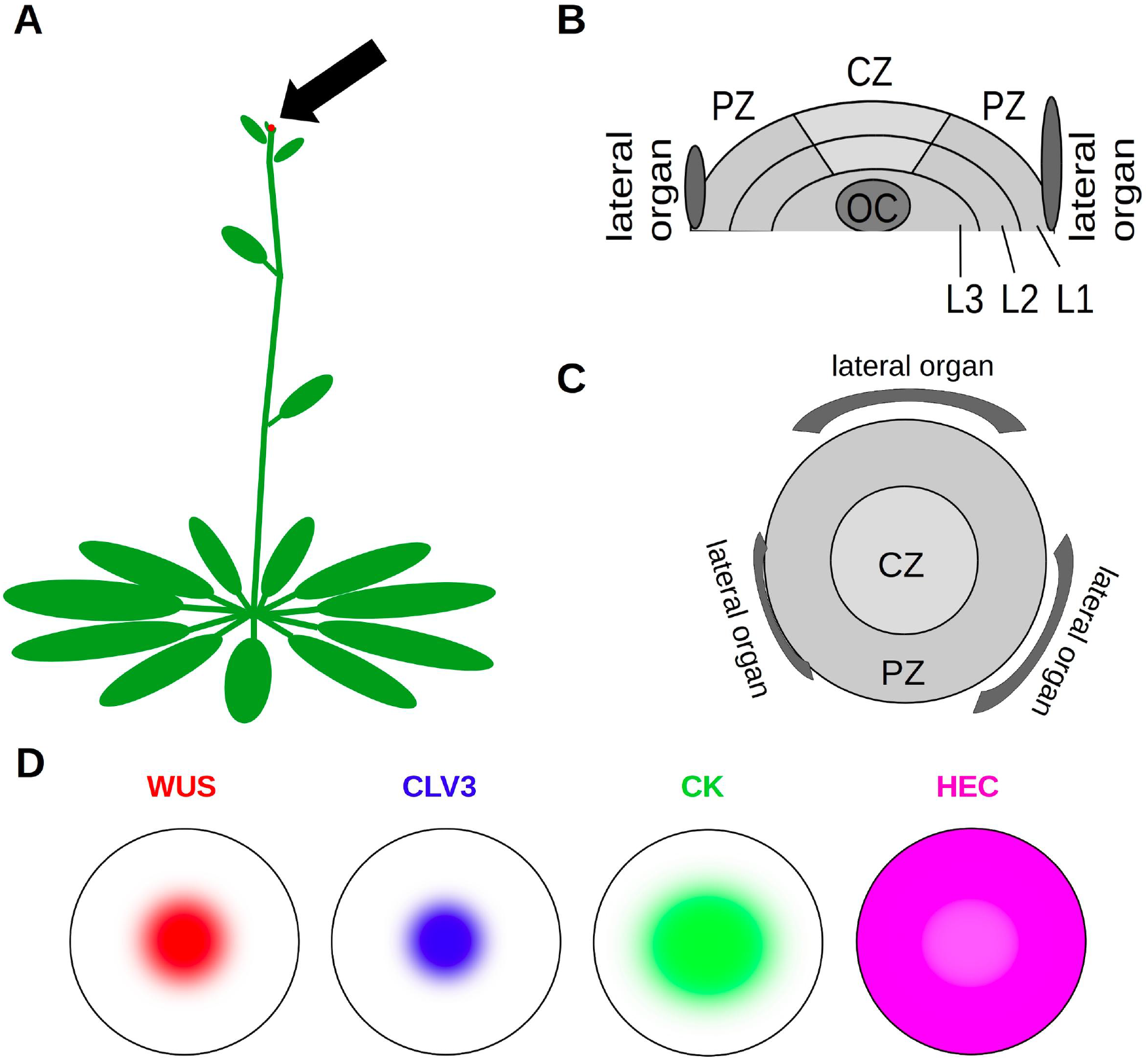
SAM (shoot apical meristem) location and morphology. (A) The SAM is located at the apex of the plant shoot (arrow). (B) Longitudinal section of the SAM. The SAM consists of three layers: L1 (epidermal cell layer), L2 (subepidermal cell layer) and L3 (corpus cell layer).(C) SAM view from the top. CZ: central zone, location of stem cells, PZ: peripheral zone, location of transit amplifying cells. OC: organizing center, location of cells producing signals that induce stem cell fate. L1, L2, L3: layers of the SAM. (D) Spatial expression patterns of key signals.

There exists experimental evidence of different transcriptional factors coordinating cell proliferation and fate choice in the SAM [6]. A key regulatory loop of the SAM consists of the transcription factors WUSCHEL (WUS) and CLAVATA3 (CLV3). WUS is produced in the OC and moves to the CZ where it maintains the stem cell identity. Stem cells in turn produce CLV3 which inhibits WUS expression in OC cells [7, 8]. This core feedback loop interacts with various other signals that fine-tune cell activity and allow the system to optimally adapt to environmental conditions [5]. One important example for such signals is the HECATE (HEC) transcription factor. It has been recently shown that HEC acts on cell fate transitions between the different SAM domains, however the underlying regulatory network remains elusive [9]. One possibility to understand the effects of perturbed signaling is the study of mutant phenotypes, such as the HEC mutant (hec123). However, studying the function of HEC genes at high spatio-temporal resolution is experimentally challenging [9]. To close this gap, we propose an integrated approach combining mathematical modeling with experimental manipulation and live imaging of plant stem cells.

Mathematical models are a powerful tool for studying complex nonlinear dynamics coordinated by multiple factors. They have contributed considerably to the understanding of SAM regulation [10–17]. In plants, cell fate decisions depend on local concentrations of spatially heterogeneous signals [5] and therefore, spatial models are required to describe meristem dynamics. For this purpose different approaches have been developed. Individual-based models allow tracking dynamics of each individual cell that is explicitly modelled. Such approach has been applied e.g., to study patterning of the WUS expressing domain [12, 16], mechanical signals [18], mechanisms of organ initiation [10], and cell fate determination [15, 19]. On the other hand, a continuous approach based on reaction-diffusion equations and ordinary differential equation models allows to study spatio-temporal interactions of different signaling molecules. Such models have been applied e.g., to investigate cytokinin signaling [11] and patterning of the shoot apical meristem [14, 17]. To investigate the impact of HEC on fate transition and proliferation rates of cells in the CZ and PZ, we have recently proposed a model based on the population dynamics approach, in which dynamics of different cell subpopulations are described by ordinary differential equations [9]. Such approach allows tracking how changes in cell proliferation, fate transition, primordia formation and primordia separation affect the time evolution and steady-state size of the different SAM zones but does not take into account spatio-temporal dynamics of the underlying signaling network.

In this paper, we study how time evolution of meristem cell populations and newly formed organs depends on the spatio-temporal dynamics of the underlying signaling network regulating cell self-renewal and differentiation. We develop a novel modeling framework that describes the SAM as a two-dimensional growing disc. The two-dimensional approximation of the domain is justified due to the SAM structure consisting of a small number of cell layers. The model describes interactions of the plant meristem key signals (CLV3, WUS, Cytokinin and HEC) and links their local concentrations to cell proliferation and fate transition rates. The change in the total SAM cell number is, in turn, linked to the change of domain size. Coupling growth and signaling processes results in a nonlinear system of reaction-diffusion equations on a growing domain with the growth velocity depending on the model components. Solving such problems is mathematically challenging. We implemented the model using the DUNE software package, which is a suitable numerical environment for sharp interface problems appearing in models with a growing domain [20–22].

The model was tested using recent experimental observations. Importantly, it allowed to gain more insight into stem cell differentiation dynamics in the HEC loss-of-function phenotype which has remained experimentally not feasible. A new insight stemming from this work is that HEC may reduce the differentiation rate of WUS producing cells [9]. The dynamics of OC cells are so far not well understood, since they are located deeply in the meristem and it is difficult to image them *in vivo*. Our model helps to understand how signals modulating the classical WUS-CLV3 loop act on the OC cells. In summary, the proposed comprehensive modeling and computational framework can be further used to generate hypotheses about interaction of the respective signaling factors and their impact on cell proliferation and fate transition.

## 1 Mathematical model and numerical computations

### 1.1 Mathematical model

#### Model geometry

Plant cells are immobile since they are encased in a cell wall. Their fate is determined by local signals; for review see [5]. The SAM consists of multiple layers. Since the cell division process is anticlinal, i.e. the progeny of cells always belong to the same layer as their parent cells [23–25], we model only the uppermost cell layer, referred to as L1, together with the organizing center (OC). Specific cross-talk signals between the L1 and L2 layer have so far not been described. Cells in both layers are exposed to the WUS signal from the OC and regulate WUS expression by production of CLV3. We model the SAM as a two-dimensional disc with radius *R*. The OC is located below the center of the meristem and it is also disc-shaped. WUS is produced in the OC and transported to the CZ [19, 26]. We model this process by a disc-shaped WUS source of radius *r* in the center of the meristem; see Fig. 2. Due to organ separation, cell proliferation and differentiation, the numbers of SAM and OC cells change over time, which results in time-dependent changes of the corresponding radii *R* and *r*.

**Figure 2:**
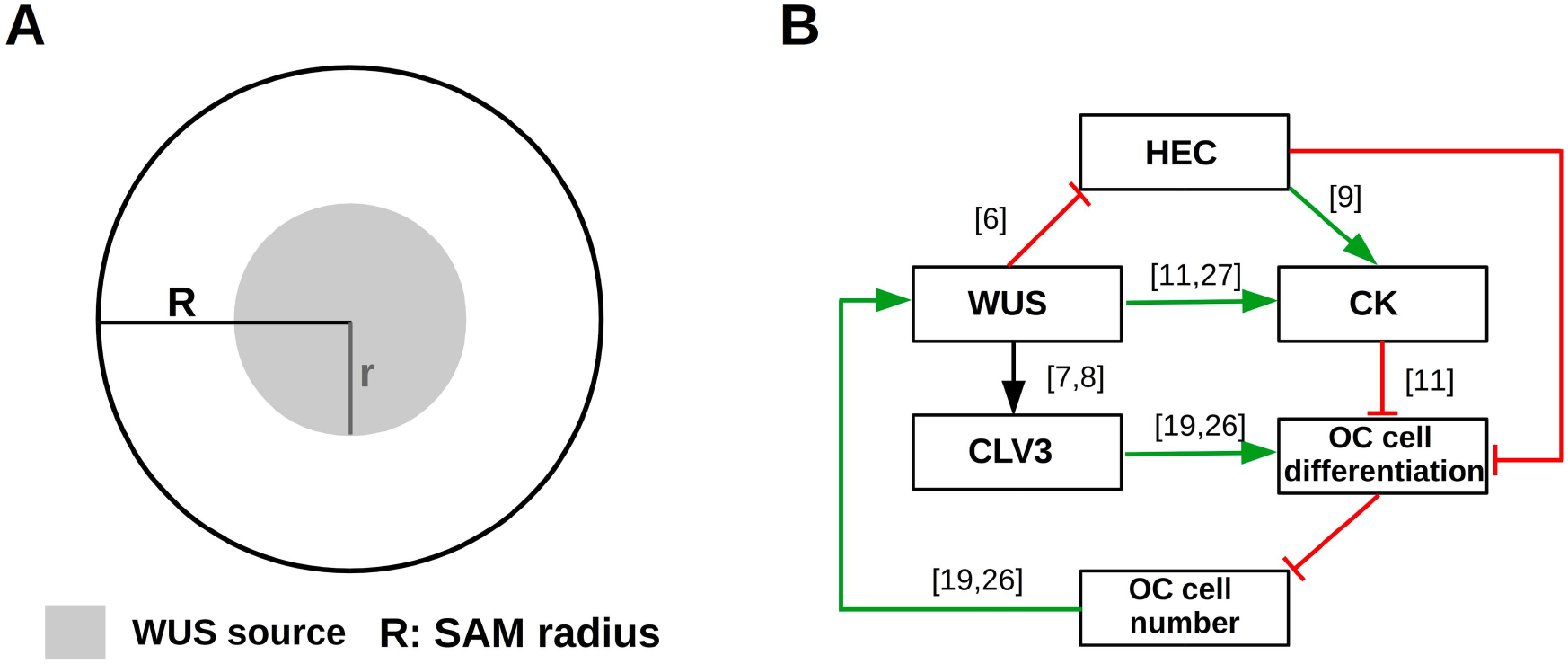
Overview of SAM geometry and regulatory feedbacks. (A) The SAM is modelled as a disc of radius *R*. The domain where WUS is produced is modelled as a concentric disc of radius *r*. (B) Regulatory signals: activating feedbacks are indicated in green, inhibiting feedbacks are indicated in red. The depicted interaction network functions at each position of the meristem. The expression domains of the respective factors evolve dynamically as a result of the interaction network, the initial condition and the diffusion of the factors. The number of OC cells determines the radius *r*. The OC is located below the SAM. The numbers in brackets correspond to the references on which the respective interaction is based.

The SAM geometry that can be represented by a spherical cap of radius *r_c_* and height *h_c_* [3] is modelled by a two-dimensional disc. The approximation results from the following procedure: The region of WUS expression is given by the radial projection of the OC on the L1 layer, which corresponds the shortest distance between the site of WUS production in the OC and the site of WUS action on L1. After the radial projection on L1, the region of WUS expression is a spherical cap of radius *r_c_* and height 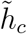. We then transform the spherical cap to a circle: Due to the rotational symmetry the spherical cap can be represented as the surface of revolution of an arc. The diameter of the circular SAM representation is equal to the length of this arc. This implies that the SAM radius equals *R* = *r_c_* cos^−1^((*r_c_* − *h_c_*)/*r_c_*), with cos^−1^ expressed in radians. Accordingly, the radius of the WUS expressing domain *r* is equal to 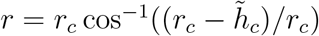.

#### Signaling network

Lateral transport of all signals is modelled by diffusion. Regulatory factors considered include WUSCHEL (WUS), CLAVATA3 (CLV3), Cytokinin (CK) and Hecate (HEC). The model accounts for the following processes:

- WUS is produced by cells in the organizing center [19, 26]. For simplicity we assume that WUS is produced at a constant rate per OC cell.
- CLV3 is produced by WUS-sensing cells [7, 8]. We assume a sigmoidal dependence of CLV3 production on WUS.
- We assume that CLV3 regulates the number of WUS-producing cells by increasing their differentiation rate or by decreasing their proliferation rate. This corresponds to a negative feedback loop between WUS and CLV3 [7, 8].
- CK signalling increases with increasing WUS concentration through inhibition of ARR5 [11,27]. Furthermore CK production increases with increasing HEC concentrations [9]. However, HEC loss of function, as in the hec123 mutant, does not completely abrogate CK production. On the other hand, WUS loss of function leads to arrest of the meristem, i.e. loss of the stem cell population [28]. We therefore assume that CK production decreases to zero in absence of WUS and that CK production is maintained at low levels in absence of HEC.
- HECATE (HEC) production is repressed by WUS [6].

In addition, we assume that all signals undergo a degradation at constant rates.

#### Description of the growing domain

We consider the Arabidopsis SAM in the inflorescence state consisting of 1600-2200 cells, corresponding to several hundreds of cells on each meristem layer [9]. The high number of cells justifies a continuous description of the SAM. In a good agreement with experimental data [9], we assume that all meristem cells have the same size. The radius of the meristem *R*(*t*) at time *t* can then be calculated based on the cell number. Let *N* (*t*) be the cell number at time *t* and *α, β* their proliferation and differentiation rates. Differentiation is defined as the commitment of meristem cells towards cells of the organ primordia: Cells located at the outer boundary of the meristem leave the meristem and contribute to the growth of the organ primordia. Cells incorporated in the organ primordia further proliferate, however their offspring do not longer contribute to SAM. Therefore, differentiation in the model leads to a decrease of meristem size. Evolution of the cell population is governed by the equation

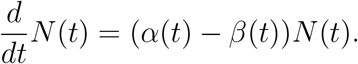

Hence, for 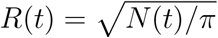, it holds 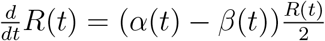.

Growth of plant structures is complex and involves multiple tightly coordinated processes [29–31], including turgor driven growth of plant cells, wall softening (hydration), wall extension, wall deposition and cell division [29, 32–34]. Our model does not resolve these processes in detail. Instead we model the change of meristem area per unit of time which corresponds to *α*(*t*) −*β*(*t*) in the above equation. This equation does not distinguish between the change in area caused by turgor-driven cell expansion alone or that involving growth and multiplication of cells.

In many of the considered scenarios, such as the hec123 phenotype or the hec over-expression phenotype, the average cell size in the SAM is constant and identical to that of the wild-type SAM [9]. In this case the cell number is strongly correlated with the SAM size.

#### Signal-dependent cell kinetics

We model the organizing center as a homogeneous cell population with signal-dependent proliferation or differentiation rate. We assume that CLV3 reduces proliferation or increases differentiation of OC cells and thus reduces the WUS concentrations. Similarly, HEC and CK reduce OC cell differentiation or induce proliferation and lead to increased WUS concentrations [11]. The considered regulatory network is summarized in Fig. 2.

We consider the following processes to describe evolution of the L1 SAM layer. WUS induces the stem cell fate [7, 8]. Stem cells have lower division rates than transit amplifying cells [9, 35]. An increase of WUS concentration leads to increase of the fraction of slowly-dividing stem cells in the meristem and decrease of cell production per unit of time. Therefore, we assume that cell production decreases with increasing total WUS concentration. As shown by experiments, reduced CK activity leads to reduction of the meristem radius [36, 37]. For this reason the growth rate of the meristem radius can become negative in presence of small CK concentrations. The mechanism underlying formation of organ primordia suggests that organ formation rates increase with the area of the meristem [10,38,39]. This implies that the cell outflux due to differentiation increases with increased meristem cell count, and hence it depends on the size of the domain and is proportional to *R*^2^. In accordance with the biological observations described above, we assume that high concentrations of CK and HEC lead to increased OC cell numbers and that high CLV3 concentrations lead to decreased OC cell numbers.

#### Mathematical model

We denote by *u*_0_(*x, t*) the concentration of WUS at location *x* and time *t*. Similarly *u*_1_(*x, t*), *u*_2_(*x, t*) and *u*_3_(*x, t*) denote the concentrations of CLV3, CK and HEC at time *t* and location *x* respectively. The meristem domain at time *t* is denoted by Ω(*t*); it corresponds to a disc of radius *R*(*t*). The organizing center at time *t* is a disc-shaped domain of radius *r*(*t*) and it is denoted as Ω_*small*_(*t*). The centers of Ω(*t*) and Ω_*small*_(*t*) coincide. The diffusion constants of WUS, CLV3, CK and HEC are denoted by *D_i_* > 0, *i* = 1, …, 4, respectively. The above-listed assumptions result in the following system of equations:

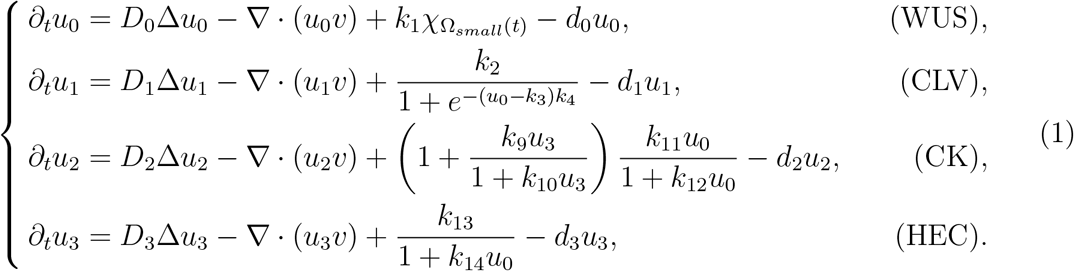

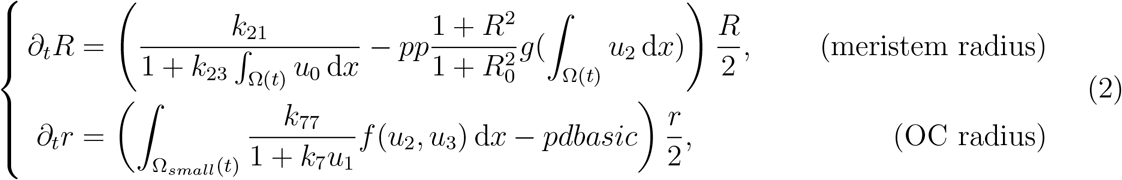

with model parameters *k_i_*, *d_i_*, *pp*, *R*_0_, *pdbasic* being positive constants. 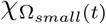 denotes the piecewise linear approximation of the indicator function of Ω_*small*_(*t*), and *v* is a function related to deformation of the domain and equal to 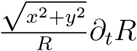. The system includes homogeneous Neumann boundary conditions in *u_i_* and initial conditions for *R*, *r* and *u_i_*. For biological reasons the radius of the organizing center cannot exceed the radius of the entire SAM. When *r* and *R* approach each other we slow down their evolution by multiplying the right hand-side of equation (2) with a nonnegative smooth function which depends on the distance between *r* and *R* and equals zero if *r* = *R*. This mechanism corresponds to physical constraints preventing that the OC has a larger diameter than the SAM.

#### Model of the domain growth

The functions *f* and *g* are defined as follows:

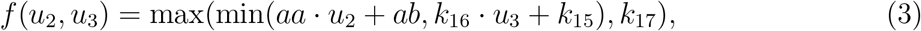

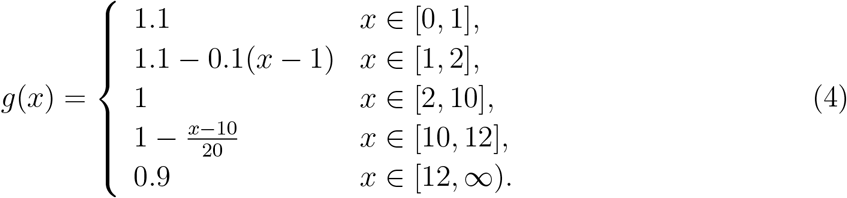

It is known that WUS expression decreases with increasing CLV3 concentrations [40]. We hypothesize that this is caused by a change of OC cell numbers. Based on experimental observations [9,11,27] we furthermore hypothesize that HEC and CK impact on OC dynamics. This agrees with experiments showing that induction of HEC in the CZ results in an increase of the OC [9]. In accordance with experimental observations showing repression of WUS by increased CLV3 concentrations [40], we consider CLV3 as the main regulator, in the sense that for high CLV3 concentrations the OC cell number decreases. We express the OC proliferation rate as the product of two functions, a decreasing Hill function depending on CLV3 and a function *f* that depends on CK and HEC. The shape of function *f* is depicted in Fig. 8 (A). Function *f* models our hypothesis that the organizing center grows if CK and HEC concentrations increase [9]. HEC is fine-tuning the meristem signaling. We assume that for increasing HEC concentrations the effect of CK on the meristem increases. This assumption follows the observations in [9]. For high HEC concentrations the CK effect saturates at a higher level compared to the case of low HEC concentrations (i.e., the impact of high levels of CK signaling is amplified by HEC). This may be explained by a HEC-induced production of CK target molecules. Since it has been observed experimentally that HEC and CK loss of function do not lead to loss of the OC [9, 37], the value of *f* is positive for *u*_2_ = *u*_3_ = 0.

Experiments have shown that decreased concentrations of CK lead to smaller meristems [36, 37]. This is modeled by the function *g* which is depicted in Fig. 8 (B) and reflects the observation that the meristem structure is robust to perturbations. Only large deviations of CK from its wild type concentration (either towards very high or very low concentrations) impact on the meristem radius by reducing the cell number [37]. The shape of the function *g* is motivated as follows. Cells in organ primordia are induced to differentiate. Since organ primordia are discrete structures, the function has multiple discrete steps. Although the organ output of the meristem decreases with decreasing CK concentrations, the number of organs per area of the meristem increases (the wildtype with a meristem diameter of 82*μm* produces 9.13 leafs within 11 days, the cre1-12 ahk2-2 ahk3-3 triple mutant with a meristem diameter of 29 *μm* produces 4 leafs within 11 days, [37]). For this reason the function *g* assumes higher values for lower CK concentrations.

It is not clear how regulation of OC size is accomplished. In principle, two extreme possibilities exist, namely constant proliferation and regulated differentiation or regulated proliferation and constant differentiation. The ODE for *r* as it is written above implies the latter. However, for uniformly bounded *u_i_* it can be rewritten as

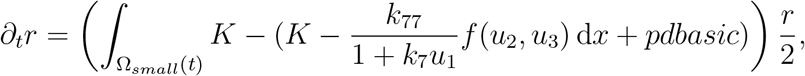

which corresponds to a constant proliferation *K* and a CLV3, HEC and CK dependent differentiation term. Here *K* denotes the maximum of 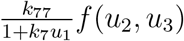. Taking into account the experimental results for the upper meristem layers [9], the first option, i.e. HEC-dependent regulation of differentiation seems more plausible.

### 1.2 Numerical approach

For numerical computations, the coupled system of reaction-diffusion equations (1) and domain evolvement (2) is decoupled using explicit equation splitting. The PDEs are then solved by the moving finite element method [41] using conforming piecewise bilinear finite elements on quadrilaterals in space and the implicit Euler method in time. The arising nonlinear algebraic system is solved with Newton’s method where the (linear) Jacobian system is solved with a sparse direct solver. The ODEs for domain movement are discretized by the explicit Euler method. Implementation has been carried out in the PDE software framework Dune/PDELab [20–22]. Details of the numerical scheme are provided in Section A11 of the Supplement.

All numerical experiments are done for mesh size *N* = 50 (i.e., 90404 degrees of freedom) and time step Δ*t* = 0.05. Following biologically relevant assumptions, we keep *r* smaller than *R*. Secondly, we take smaller time steps if radii change very rapidly. One of main problems in numerical simulation is the form of the evolutionary equation describing the dynamics of *r*. To calculate its right-hand side, we integrate a nonlinear function which depends on the model solution. It poses a numerical error. Hence, estimating parameters *pdbasic* and *pp* based on the prescribed steady-state values of *r* and *R* leads to different parameter values depending on the mesh size. The values of *pp* and *pdbasic* used in numerical analysis of the model are values obtained in a limiting procedure. That explains why in Fig. 3 (C) we obtain the different values of inner radius for different mesh sizes.

**Figure 3:**
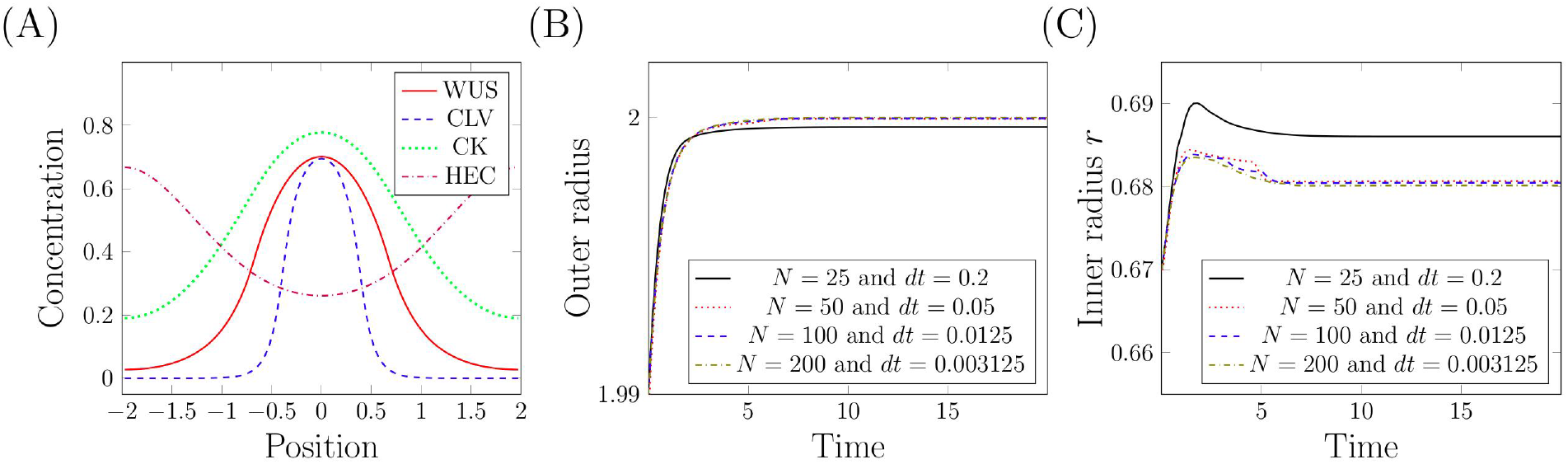
Numerical calculation of the steady-state. (A) For the depicted initial condition the system approaches the equilibrium depicted in Fig. 6 (A). The corresponding evolution of *R* and *r* is depicted in (B) and (C) respectively. Calculations have been performed for different mesh sizes *N* and time steps Δ*t*: *N* = 25 and Δ*t* = 0.2 (22704 degrees of freedom), *N* = 50 and Δ*t* = 0.05 (90404 df), *N* = 100 and Δ*t* = 0.0125 (360804) and *N* = 200 and Δ*t* = 0.003125 (1441604 df).

### 1.3 Model calibration

#### Initial data and model parameters

Since stem cells are identified experimentally using CLV3 reporters, we define them in the model as the cells located at positions where CLV3 concentration is above a certain threshold. We assume that the meristem of the unperturbed adult wild type plant is in a steady-state. Experiments show that under such conditions the CZ cell number corresponds to approximately 10% of the total meristem cell number [9]. This ratio is hold in the locally stable equilibrium depicted in Fig. 6 (A). This equilibrium serves as a departure point for all simulated experiments. The corresponding model parameters are listed in Table 1. These provide an example set of parameters that fit the wild type meristem configuration. Details on the calibration of proliferation and differentiation rates are given in Section A10 of the Supplement. A sensitivity analysis is performed in Section A12 of the Supplement.

**Table 1:** Parameter values corresponding to the wild type (unperturbed) scenario.

| name | value |  | name | value |  | name | value |  | name | value |
| --- | --- | --- | --- | --- | --- | --- | --- | --- | --- | --- |
| $D_0$ | 1 | | $D_1$ | 1 | | $D_2$ | 0.2 | | $D_3$ | 0.2 |
| $d_0$ | 5 | | $d_1$ | 64 | | $d_2$ | 1.1 | | $d_3$ | 0.6 |
| $k_1$ | 6 | | $k_2$ | 50 | | $k_3$ | 0.6 | | $k_4$ | 100 |
| $k_9$ | 4 | | $k_{10}$ | 0.1 | | $k_{11}$ | 1.4 | | $k_{12}$ | 1 |
| $k_{13}$ | 2 | | $k_{14}$ | 100 | | | | | | |
| $k_{21}$ | 1.65 | | $k_{23}$ | 0.01 | | $pp$ | 1.622 | | | |
| $k_7$ | 500 | | $k_{77}$ | 20 | | $pdbasic$ | 0.83 | | | |
| $aa$ | 2 | | $ab$ | 0.4 | | $k_{15}$ | 0.4 | | $k_{16}$ | 5 |
| $k_{17}$ | 1.1 | | | | | | | | | |

**Table 2:**
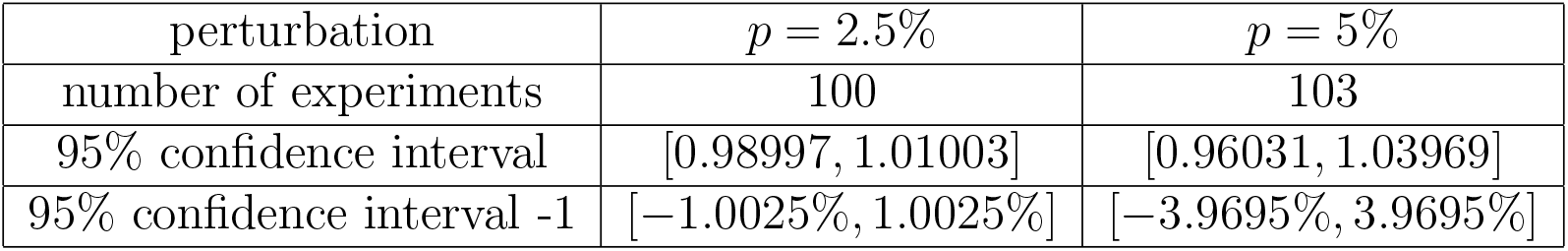
Changes of the ratio *r/R* for randomly perturbed model parameters. Changes of the ratio *r/R* if model parameters are randomly perturbed with values from the interval [−*p, p*]. For *p* ∈ {2.5%, 5%} at least 100 numerical experiments were simulated. The numerical value *r/R* assumes in the system with unperturbed parameters simulated on the same mesh with the same time step corresponds to 100%. It equals 0.3403.

| perturbation | $p = 2.5\%$ | $p = 5\%$ |
| --- | --- | --- |
| number of experiments | 100 | 103 |
| 95% confidence interval | [0.98997, 1.01003] | [0.96031, 1.03969] |
| 95% confidence interval -1 | [-1.0025%, 1.0025%] | [-3.9695%, 3.9695%] |

#### Stationary state

The stationary state which serves as departure point for the simulated experiments has been found numerically. The initial condition used to converge to this equilibrium is depicted in Fig. 3 (A), the time evolution of *R* and *r* in Fig. 3 (B), (C). The choice of the steady state as the initial condition is supported by experiments relying on inducible changes of gene expression. The plant is grown to the inflorescence state and only then the changes in gene expression are induced using e.g., dexamethasone or ethanol [9, 40, 42–44]. Control measurements demonstrate that in the absence of the induced changes the meristem size remains constant and is identical as in the wild type.

In the following section, we provide mathematical evidence for the local stability of this steady state.

For this formal reasoning, we assume that the reaction-diffusion process of signaling molecules is faster than changes of the domain radius, we obtain a quasi-stationary system. For a given pair (*R, r*) we solve the quasi-stationary problem for ***u***, i.e. setting the time derivatives in (1) equal to 0 we obtain the solution ***u*** as a function of *R* and *r*. Note that the (WUS) equation of system (1) for given (*R, r*) is a linear elliptic equation that can be solved explicitly. It allows further obtaining *u*_1_ and *u*_3_ from the (CLV) and (HEC) equations, respectively, and finally *u*_2_ from the (CK) equation. Inserting the obtained solution ***u*** into Eq. (2) provides an evolutionary system for (*R, r*):

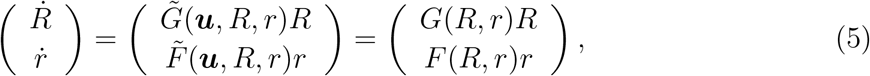

where functions 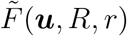 and 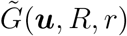 correspond to the expressions on the right-hand of system (2), see Fig. 4. Linearization in the neighborhood of the steady-state (*R*\*, *r*\*) leads to

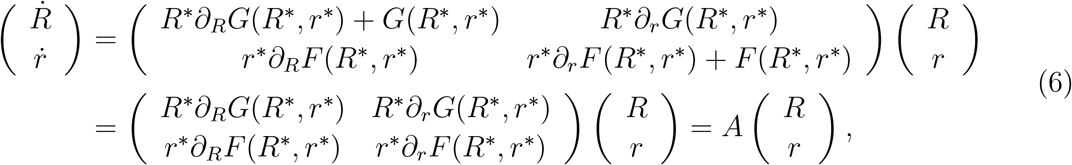

since *F* (*R*\*, *r*\*) = 0 and *G*(*R*\*, *r*\*) = 0.

Derivatives of *G* are negative, what can be checked by explicit calculations. The main task is to calculate derivatives of function *F*. We are not able to do it analytically. However, using our numerical approach we can calculate values of the function *F* in the neighborhood of the steady-state, see Fig. 5. Hence, we obtain that *∂*_*r*_*F* is negative and *∂*_*R*_*F* is positive.

**Figure 4:**
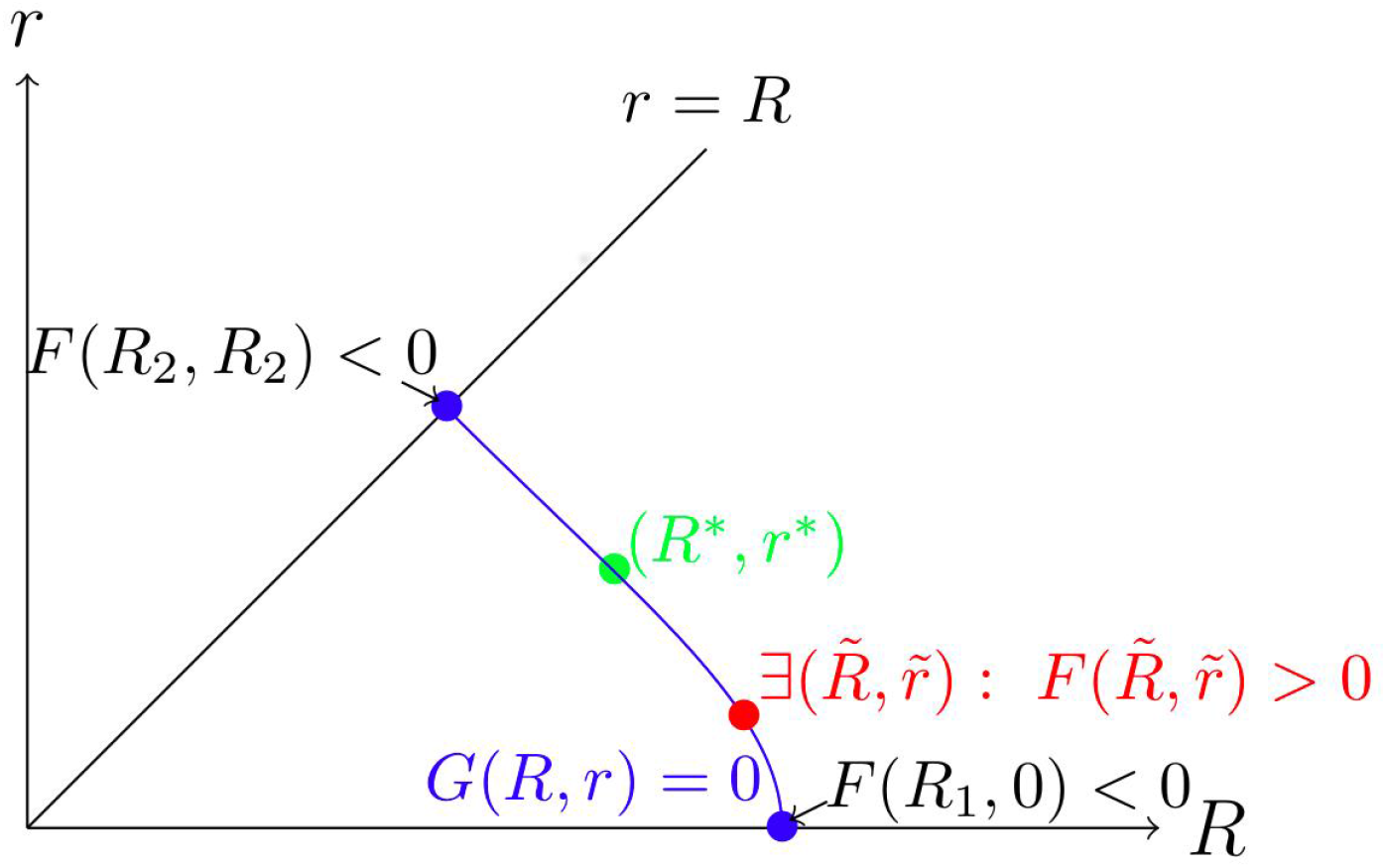
Existence of steady state. The blue curve describes the zero level-set of function The points (*R*_1_, 0) and (*R*_2_, *R*_2_) correspond to the intersection of the zero level-set of function *G* with the lines *r* = 0 and *r* = *R*. The value of the function *F* in these points is negative. For *r* = 0 and *r* = *R* we are able to solve (1) explicitly. Moreover, we know that there exists at least one point 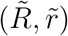 for which function *F* is positive. The latter is a consequence of the parameter choice. Thus there exists at least one point (*R*\*, *r*\*) such that: *G*(*R*\*, *r*\*) = 0, *F* (*R*\*, *r*\*) = 0 and in the neighborhood of this point for *R* > *R*\* it holds *F* (*R*, *r*\*) ≥ 0 and for *R* < *R*\* it holds *F* (*R*, *r*\*) < 0. Further on, we will consider the stability of the steady state solution (*R*\*, *r*\*).

**Figure 5:**
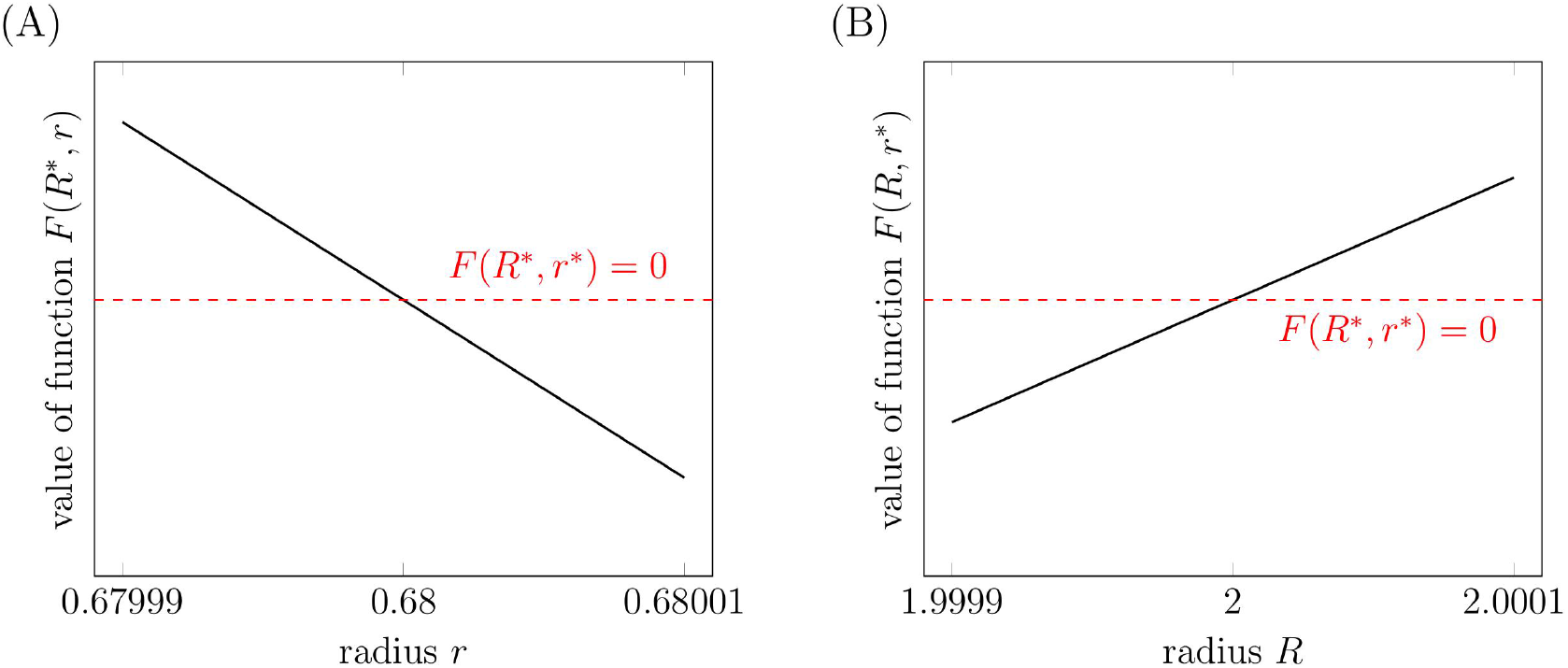
Local stability of the steady state. Both plots present values of the function *F* (*R, r*). In the left panel we set *R* = *R*\*, in the right panel we set *r* = *r*\*. The dashed red lines correspond to *F* (*R*\*, *r*\*) = 0. All calculations were done for mesh size *N* = 50 (90404 degrees of freedom). Also the value *F* (*R*\*, *r*\*) is calculated using the value of *pdbasic* corresponding to the mesh size *N* = 50.

Consequently,

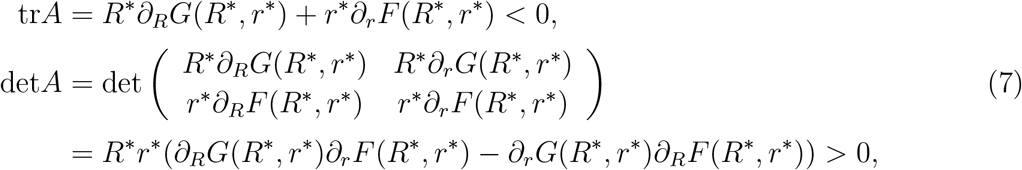

since *r*\* and *R*\* are positive. This implies that the steady-state is locally stable. We assumed the quasi-steady state for the formal analysis only. Notably, all simulations shown consider the fully coupled system without any quasi-steady state reduction.

#### Experiments used to test the model

The model is tested by comparing its results to the following experiments.

- WUS over-expression in the whole meristem: This leads to radial expansion of the CLV3 expressing domain. The change of total SAM size is negligible [42].
- WUS loss of function: This leads to termination of the meristem and loss of stem cells [28].
- CLV3 over-expression in the central zone: This leads to repression of WUS, it has been shown that a 10 fold change in CLV3-expression levels does not affect meristem size [40].
- CLV3 loss of function: This leads to larger meristems and expansion of the CZ [44].
- Reduced degradation of CK: This leads to larger meristems and a larger OC [45].
- CK loss of function: This leads to significantly smaller meristems [36, 37].
- HEC over-expression by stem cells: This leads to expansion of the central zone and subsequent loss of meristem structure [9].
- HEC loss of function: The HEC triple mutant hec123 expresses no functional HEC. This leads to significantly smaller meristems compared to the wild type [9].

## 2 Results

### 2.1 Simulation of key experiments

In this section, we test the model by comparing it to the outcomes of a set of experiments involving over-expression or loss-of-function of certain signals. In several places, we refer to genes expressed under a promoter. To express gene X under the promoter of gene Y means to engineer genes such that X is always expressed together with Y. If the promoter of Y is ubiquitously expressed, then X is expressed everywhere in the meristem, if the promoter of Y is site specific, then X is expressed only in a subdomain of the meristem.

#### 2.1.1 Perturbation of WUS

##### WUS over-expression

There exist different experimental works studying an ubiquituous increase of WUS [42, 43]. Experimentally this has been accomplished using a glucocorticoid-inducible form of WUS under a promoter that causes ubiquitous expression. The experimental setting is modelled by the following modification of the equation for WUS:

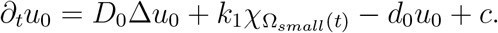

The positive constant *c* denotes the rate of WUS over-expression which is independent of space, time and other signals. In the considered plants the induced ubiquitous WUS expression acts in addition to the physiological WUS expression in the organizing center. Therefore, the equation contains both 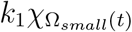 and *c* source terms. The steady-state shown in Fig. 6 (A) serves as initial condition for simulation of the experiment.

**Figure 6:**
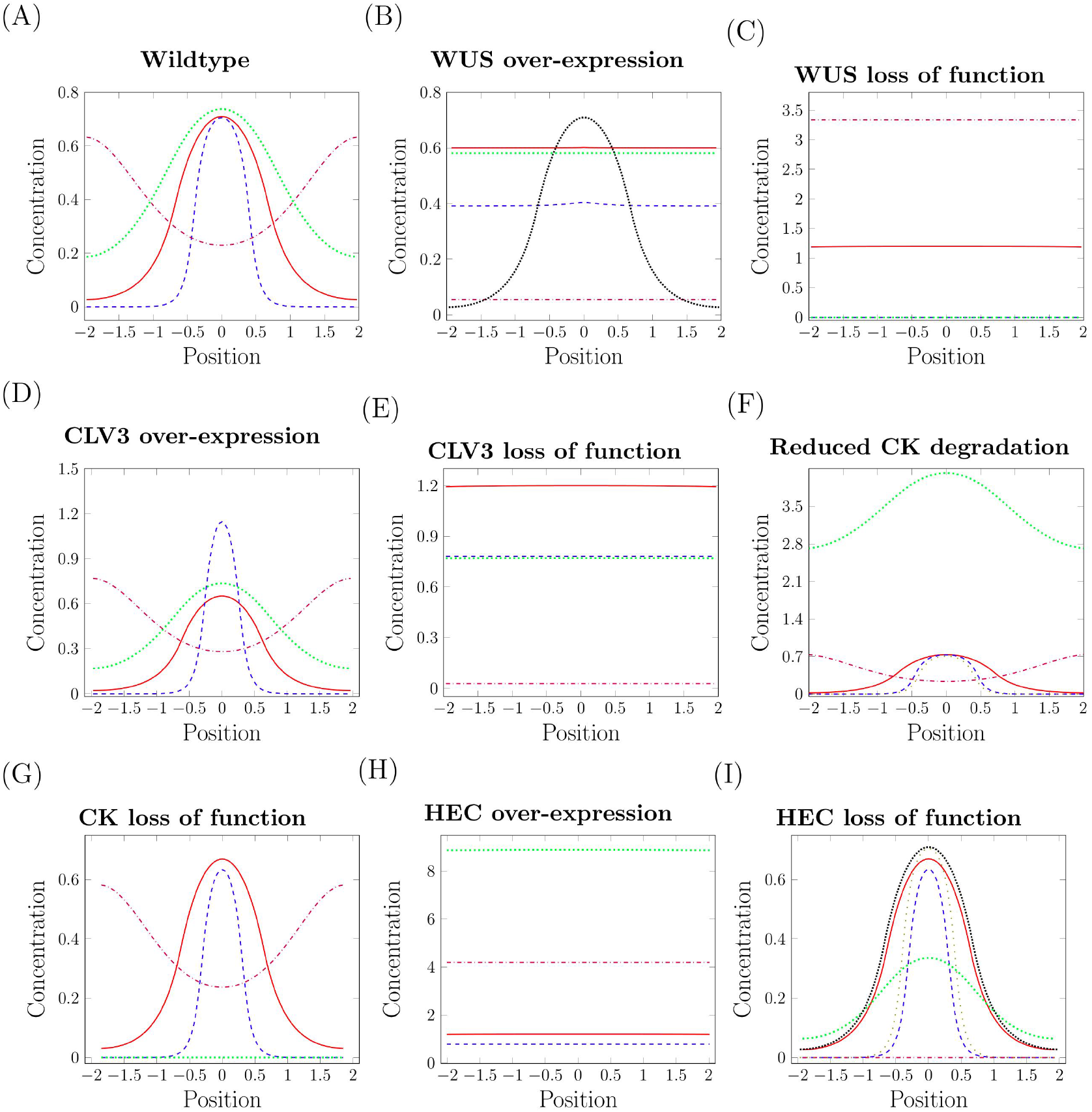
Signal concentrations in over-expression and loss-of-function experiments. Signal concentrations along the diameter of the meristem, position 0 corresponds to the center: WUS (solid red), CLV3 (dashed blue), CK (dotted green) and HEC (dashdotted purple). (A) Unperturbed steady-state of the wild-type meristem. (B) Ubiquitous over-expression of WUS: The system converges to a state with CLV3 expression in the whole meristem. The equilibrium concentration of WUS in the wild-type is shown for comparison (densely dotted black). (C) WUS loss of function: The system converges to a steady-state with negligible CLV3 concentrations, which corresponds to the experimentally observed loss of stem cells. Breakdown of the negative feedback between WUS and CLV3 leads to high concentrations of WUS molecules that are not functional. (D) CLV3 over-expression: The system converges to a state with higher CLV3 and slightly reduced WUS concentrations. (E) CLV3 loss of function: Due to the missing negative feedback the OC expands and the CZ spreads over the whole meristem. (F) Reduced degradation of CK: The system converges to a state with constant in space signal concentrations. For comparison the CLV3 profile of the wild-type steady-state is depicted (loosely dotted olive). (G) CK loss of function does not lead to significant changes in the simulations. (H) HEC over-expression in stem cells: The central zone expands until it reaches the boundary of the meristem. (I) HEC loss of function (so called HEC triple mutant): The system converges to a state with lower WUS and CLV3 concentrations and a smaller meristem. HEC concentration (dashdotted purple) is equal to zero. For comparison, the CLV3 (loosely dotted olive) and WUS (densely dotted black) profiles of the wild-type steady-state are depicted.

The biological experiments agree in the observation that the central zone gets larger. This we also observe in the simulations. If the over-expression is high enough, the simulations show a radial expansion of the CLV3 expression domain and the system converges to a state where CLV3 is expressed in the whole meristem; Fig. 6 (B). This observation agrees with the experimental results from [42]. The radial expansion of the CLV3 expressing domain is depicted in Fig. 10. Biologically it has been considered unexpected that ubiquitous WUS over-expression leads to a radial growth of the CZ instead of a simultaneous up-regulation of CZ fate in the whole meristem. It has been speculated that the reason for this observation is that PZ cells located at the boundary of the CZ respond differently to WUS compared to other PZ cells [42]. The model simulations, however, suggest that the experimental observations can be explained even if all PZ cells respond to WUS equally. The observation that the total meristem size does not change significantly in the simulation matches also the experimental findings [42]. In the simulations we see that ubiquitous WUS expression is linked to a reduction in proliferation rate, see Figure 18 (B), as it has been observed in experiments [42].

Simulations predict different outcomes for different levels of over-expression, Fig. 9 in Supplement. Since it is experimentally difficult to fine-tune the rate of over-expression, the diverse outcomes obtained in the simulations have not yet been observed.

##### WUS loss-of-function

To simulate WUS loss of function, we set the WUS concentration equal to zero in the equations for CLV3, HEC and CK. In this setting WUS is still produced but it does not impact on dynamics of other signals. This corresponds e.g., to the case where WUS cannot bind to its receptors. This scenario results in loss of stem cells, since the defective WUS protein cannot induce production of CLV3. Experimentally such a scenario can be studied using WUS loss-of-function mutants, as it has been done in [28] or inducible RNAi [26,42,43]. The loss of stem cells observed in the simulations is in agreement with experimental data. Due to the low CLV3 levels, there exists no feedback inhibition of WUS expression which implies that levels of the non-functional WUS protein are high and the organizing center grows. For this reason the system converges to a state with negligible CLV3 concentrations and high WUS expression as shown in Fig. 6 (C). Time evolution of *R* and *r* is depicted in Fig. 11. As in the experiments, WUS loss of function leads to the loss of the stem cell population which is characterized by CLV3 expression and slow division [28].

#### 2.1.2 Perturbation of CLV3

We simulate a scenario where CLV3 expression is increased in its natural expression domain. This has been experimentally accomplished in [40] by using an ethanol-inducible CLV3 construct that is expressed under the CLV3 promoter. We implement this by multiplying the CLV3 production term by a positive constant *c*:

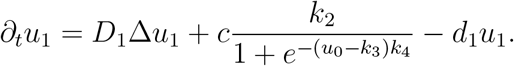

Model simulations predict the outcome of experiments showing only mild changes in total meristem size. We also obtain reduced WUS concentrations. Such dynamics have been reported as a transient phenomenon in experiments [40]. Simulation results are shown in Fig. 6 (D) and Fig. 12.

We implement CLV3 loss-of-function by setting CLV3 concentrations to zero in the right hand-side of the equations for WUS, HEC, CK, *r* and *R*. Experimentally such a scenario has been accomplished through CLV3 silencing using inducible RNA interference [44]. As in experiments, we observe expansion of the central meristem zone [44] and an increased proliferation rate in the center [46]. However, we do not observe the reported expansion of the total SAM size. This suggests that there exists a coupling between CZ size and PZ proliferation rate that is not considered in the model and has not yet been characterized in detail. Results are shown in Fig. 6 (E), Fig. 18 (E) and Fig. 13.

#### 2.1.3 Perturbation of CK

We simulate the following scenarios of CK perturbation:

- CK over-expression: during this experiment we change the degradation rate in the equation for CK;
- CK loss of function: during this experiment we put 0 instead of the production term in the CK-equation.

To study the impact of increased CK concentration the ckx3 ckx5 double mutant has been used. In this mutant the degradation of CK via CKXs (cytokinin oxigenases/dehydrogenases) is reduced [45]. We model this experiment by reducing the value of *d*_2_, which corresponds to decreased CK degradation, as in the experiments. Numerical simulations are consistent with the experiments in showing an increase of the OC, a slight increase of the CZ and an increase in meristem radius [45].

Arabidopsis mutants lacking functional CK receptors, such as the cre1-12 ahk2-2 ahk3-3 triple mutant allow to study CK loss of function [37]. In agreement with experiments [36,37] simulations show a decrease of the OC radius and of the total meristem size, see Figure 15.

### 2.2 New insights arising from *in silico* perturbation of HEC signaling

#### 2.2.1 Simulation of HEC overexpression and loss-of-function

Recently, we have established an experimental setting to study HEC over-expression in stem cells using an inducible HEC1 form expressed under the CLV3 promoter. The experiments show a change of meristem size and stem cell number in case of HEC over-expression [9]. Our hypothesis is that HEC acts on the OC cell differentiation or proliferation. This cannot be directly monitored in experiments but can be tested using our model. We implement this experiment in the model by adding a HEC production term that is proportional to the CLV3 source term in equation (1). This yields the following equation for HEC:

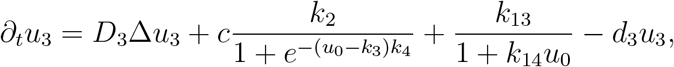

where *c* describes the proportionality between HEC and CLV3 production resulting from the expression of both signals under the same promoter. This scenario corresponds to the experimental setting from [9], where HEC is expressed in stem cells that are characterized by high CLV3 levels. For *c* large enough we observe that the system converges to a state with constant in space signal concentrations. This corresponds to the experimentally observed expansion of the CZ towards the boundaries of the meristem. Identically as in the experiments WUS, CLV3 and CK concentrations are increased compared to the wild-type meristem, Fig. 6 (H) for *c* = 3. In the simulations, see Fig. 18 (H), as in the experiments [9], the proliferation rate is reduced in the central zone and increased at the boundary of the meristem compared to the wild-type. In agreement with experiments we observe an increase in total meristem radius *R*; Fig. 16.

We implement HEC loss of function by setting HEC production equal to zero. As in experimental data [11], we observe mild changes in WUS and CLV (Fig. 6 (I)) that are associated with a smaller meristem (Fig. 17). The flux of cells from the meristem into lateral organs is increased in the simulations compared to the wild-type. This is in agreement with the results from [9]. Furthermore we see only very slight changes in proliferation rates which also fits to the conclusions from [9].

#### 2.2.2 Model simulations suggest a direct effect of HEC on OC cells

In Fig. 2 (B) we propose a direct action of HEC on OC cell differentiation and an indirect action via CK. If we repeat simulation of the experiment omitting the direct action of HEC on OC cells, we cannot reproduce the wet lab experiment, since in the simulation the CZ does not extend towards the boundaries of the meristem Fig. 7. This observation supports the existence of a direct effect of HEC on OC cells.

**Figure 7:**
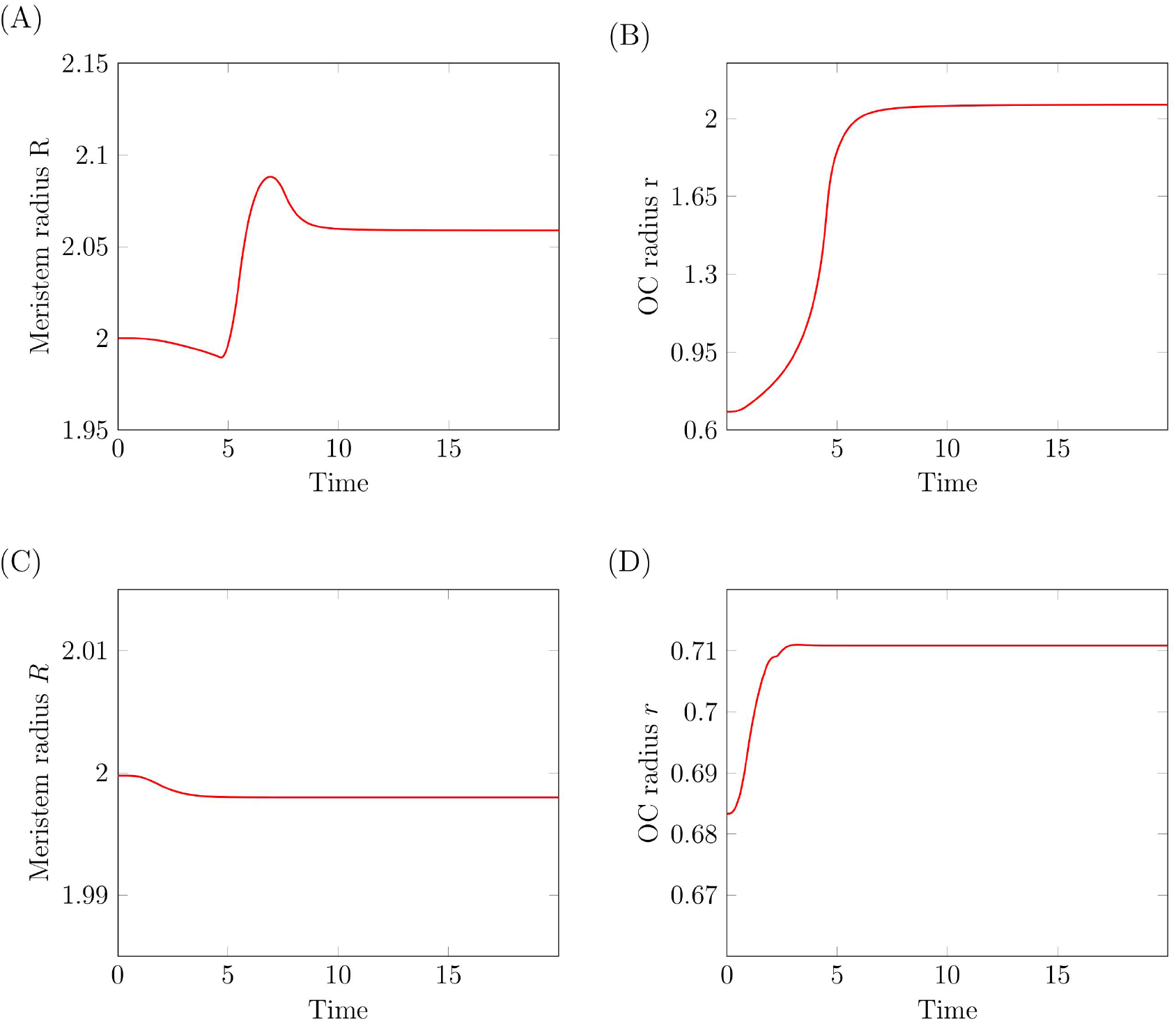
Direct effect of HEC on OC cells. Simulation of HEC over-expression with (A-B) and without (C-D) a direct effect of HEC on OC cell differentiation. (A-B) Time evolution of meristem and OC radius during HEC over-expression in stem cells. The simulations assume a direct effect of HEC on OC cells. (C-D) Time evolution of meristem and OC radius during HEC over-expression in stem cells. The simulations assume no direct effect of HEC on OC cells. Unlike in the experiments the WUS expressing domain does not extend over the whole meristem. For all depicted simulation we set *c* = 0.05.

## 3 Discussion

Plant shoot apical meristem systems are tightly regulated. However, the question how gene regulatory networks control the transition from stem cells to fully differentiated cells located in lateral organs is still open. There is evidence that the key negative feedback loop consisting of WUS and CLV3 is modulated by other hormonal signals. Recently it has been established that HEC is an important regulator of cell fate transition in the shoot apical meristem. Experimental works have studied the interaction of HEC with relevant plant hormones and its impact on cell properties [5, 6, 9].

The model framework developed in this work allows to consider the interaction between different regulatory signals and their impact on cell kinetics and fates. The 2d geometry accounts for the planar heterogeneity of signal expression. The growing domain framework keeping track of changes in SAM cell number allows to test hypotheses that relate signal concentrations to cell proliferation and differentiation.

The model developed in this work is based on the assumption that HEC inhibits the differentiation (or promotes the proliferation) of OC cells leading to an increase of the WUS expressing domain and thus to an increase of the CZ. We simulate the impact of the configuration of the regulatory network on the time evolution of signal concentrations, OC size, CZ size and total meristem cell count. Using fluorescence constructs all these quantities are experimentally accessible. The proposed regulatory feedbacks are able to qualitatively reproduce meristem changes under WUS loss of function, CLV3 loss of function, HEC over-expression, HEC loss of function and CK loss of function. This supports the hypothesis that HEC directly and indirectly (via CK) leads to a reduction of OC cell differentiation (or an increase of OC cell proliferation).

Our simulations lead to several new biological insights. It is known that HEC is not expressed in OC cells and that experimental expression of HEC in the OC leads to the loss of the meristem [6]. Our model suggests that HEC directly acts on OC cell kinetics and that this action is required to observe the growth of the CZ in HEC over-expression experiments.

In addition to this there exists a CK mediated effect of HEC on OC cell differentiation: HEC increases CK expression and CK affects OC cells. This HEC mediated increase of CK signaling is required to explain the reduced CK levels observed in the hec1,2,3 triple mutant [9]. The effect of CK on OC cell differentiation explains the change of OC size under CK perturbation. Together these control couplings constitute an indirect CK-mediated effect of HEC on OC cell differentiation. It is difficult to predict intuitively whether this indirect effect is sufficient to explain experimental results or whether an additional direct action of HEC on the OC is required. Our simulations suggest that without the direct effect the WUS expressing domain does not show the gradual increase until it reaches the boundary of the meristem as it has been observed experimentally in [9]. Therefore, our simulation support the existence of a direct effect of HEC on OC cell kinetics. The investigation of HEC target genes may help to identify reguatory nodes mediating the effect of HEC and to integrate them with known candidates such as NGATHA [47].

In case of CLV3 loss of function or increased CK activity, the model predicts an increase of the CZ what is reflected by experiments [44, 45], however it fails to reproduce the observed growth of the meristem. The reason for this is that in the model system PZ cells are recommitted to the stem cell fate, due to increase of WUS activity, which leads to an increase of the pool of slowly dividing stem cells at the expense of fast dividing PZ cells. Hence, the model results in disappearance of the PZ. The discrepancy between the model predictions and the experimental data showing maintenance of the PZ suggests that there exists an additional mechanism compensating the loss of PZ due to loss of CLV3 function. A molecular mechanism underlying this observation is not known. In [44], the authors observe an increase of PZ cell proliferation rates in case of CZ expansion and refer to it as a “long-distance effect”. Similarly, in [9] an increase in the PZ mitotic index is observed following the CZ expansion due to HEC over-expression in stem cells. Taking into account that HEC is unable to move from cell to cell, the latter observation suggests that there exists a non-cell autonomous mechanism that adjusts the number of PZ cells to the size of the CZ. Our model verifies that CK signalling is not sufficient to mediate this effect. In [42] it is discussed that a dose-dependent inhibitory effect of WUS on proliferation rates may play a role in this context. Furthermore, ERECTA signalling may be involved in confining the WUS and CLV3 expression to the meristem centre by repressing both signals [48, 49]. These mechanisms could be compared in future extension of our model.

Future version of the model have to include auxin signaling. It is known that auxin promotes cell fate transition from the PZ to lateral organs. Auxin signaling is repressed by WUS [43] in the meristem center and modulated by HEC in the meristem periphery [9]. The proposed computational framework is ideally suited to investigate details of these interactions.

In conclusion, we have developed a modeling framework that allows to study how gene regulatory networks control fate transition dynamics in the shoot apical meristem. We propose a network configuration that is sufficient to reproduce key experiments. Our simulations provide new insights in the effect of HEC and lead to the hypothesis that HEC directly and indirectly (via CK) reduces OC cell differentiation rates and thus induces an expansion of the central zone of the meristem if it is over-expressed in stem cells.

## Appendix A1. Functions *f* and *g*

The shapes of the functions *f* and *g* are depicted in Fig. 8.

**Figure 8:**
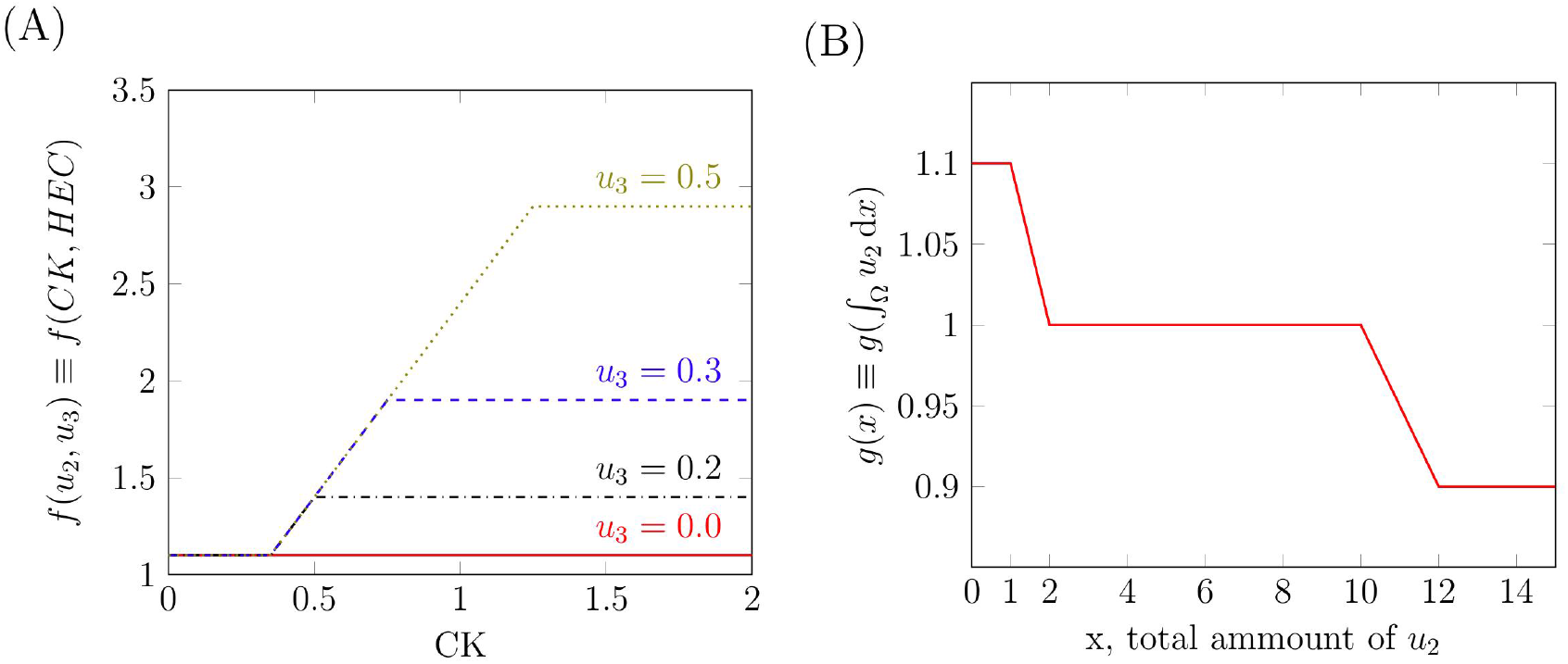
Functions *f* and *g*. (A) The shape of the function *f* for different concentrations of CK and HEC. (B) The shape of the function *g*.

## Appendix A2. Different rates of WUS over-expression

We run simulations for different rates of WUS over-expression. In the model this corresponds to different values of *c*. Depending on the rate of WUS over-expression different dynamics can be observed.

- Mild WUS over-expression (*c* = 1.0) in the whole meristem, Fig. 9 (A): Inhibition of HEC leads to increased OC differentiation. The OC shrinks and therefore also the WUS and CLV3 concentration in the center of the meristem. The over-expression is too mild to trigger a CLV3 production comparable to that in the center of the wild type CZ. The system approaches a state with decreased WUS and CLV3 concentrations. The OC cell number is different from zero and the state is nonconstant in space.
- Intermediate WUS over-expression (*c* = 2.9) in the whole meristem, Fig. 9 (B): Inhibition of HEC leads to increased OC differentiation and extinction of the OC cells. Therefore, WUS production in the OC ceases. The system converges to a state that is constant in space and that is maintained by the ubiquitous constant WUS expression. This level of WUS expression, however, leads to CLV3 concentrations lower than the CLV3 concentrations in the center of the wild-type meristem.
- Strong WUS over-expression (*c* = 3.1) in the whole meristem, Fig. 9 (C): Inhibition of HEC leads to increased OC differentiation and extinction of OC cells. The system converges to a state that is constant in space and that is maintained by the ubiquitous constant WUS expression. In this case the WUS expression is high enough to trigger CLV3 concentrations similar to those in the center of the wild-type meristem. This case reflects what is seen in WUS over-expression experiments [42].

**Figure 9:**
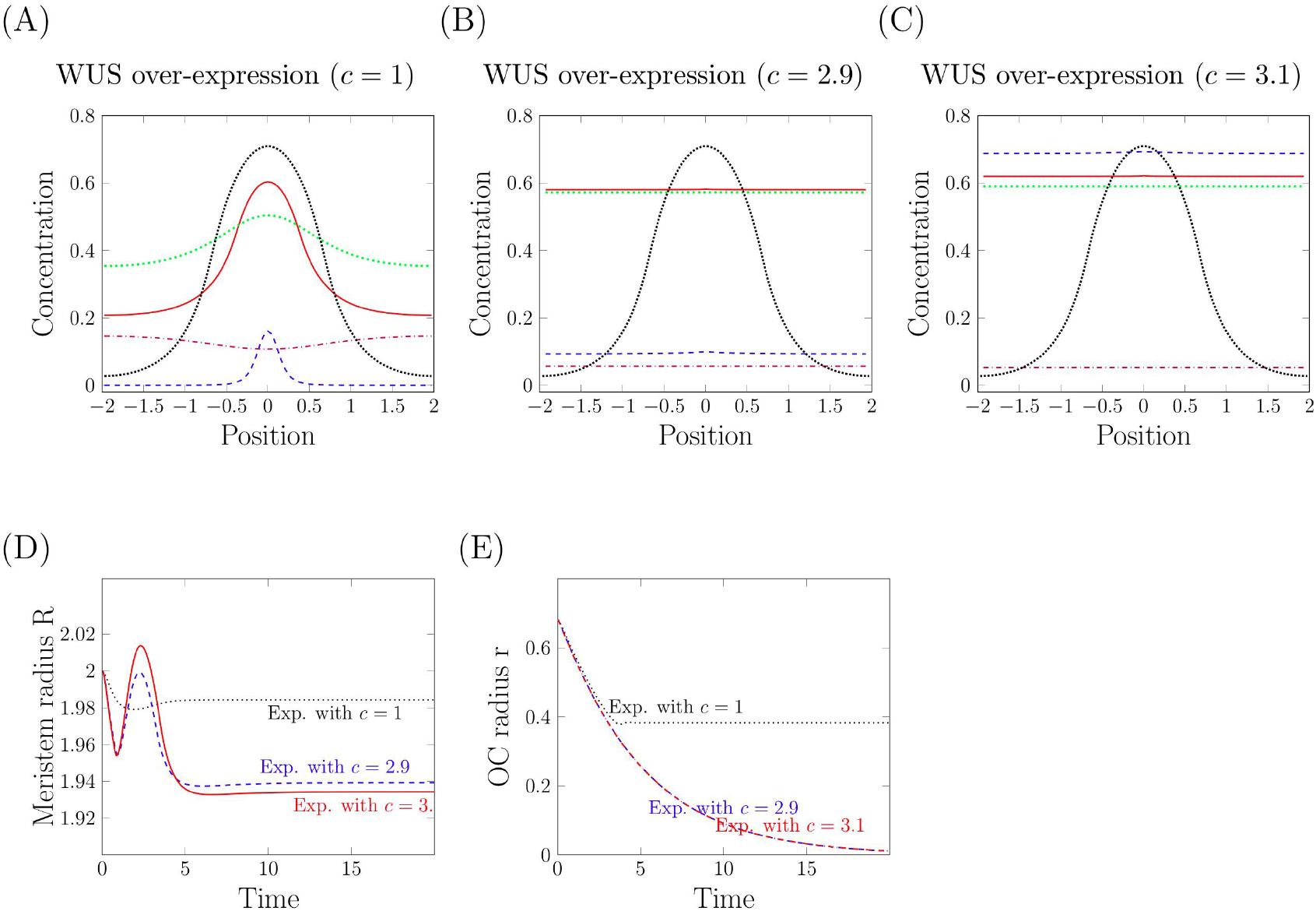
Mild, intermediate and strong WUS over-expression. Concentrations of signals along the diameter of the meristem, position 0 corresponds to the center: WUS (solid red), CLV3 (dashed blue), CK (dotted green) and HEC (dashdotted purple). WUS over-expression is simulated by adding a constant production *c* in the whole meristem. (A) Mild ubiquitous WUS over-expression. (B) Intermediate ubiquitous WUS over-expression. (C) Strong ubiquitous WUS over-expression. The equilibrium concentration of WUS in the wild type is shown for comparison (densely dotted black). (D, E) Time evolution of meristem and organizing center (OC) radius.

**Figure 10:**
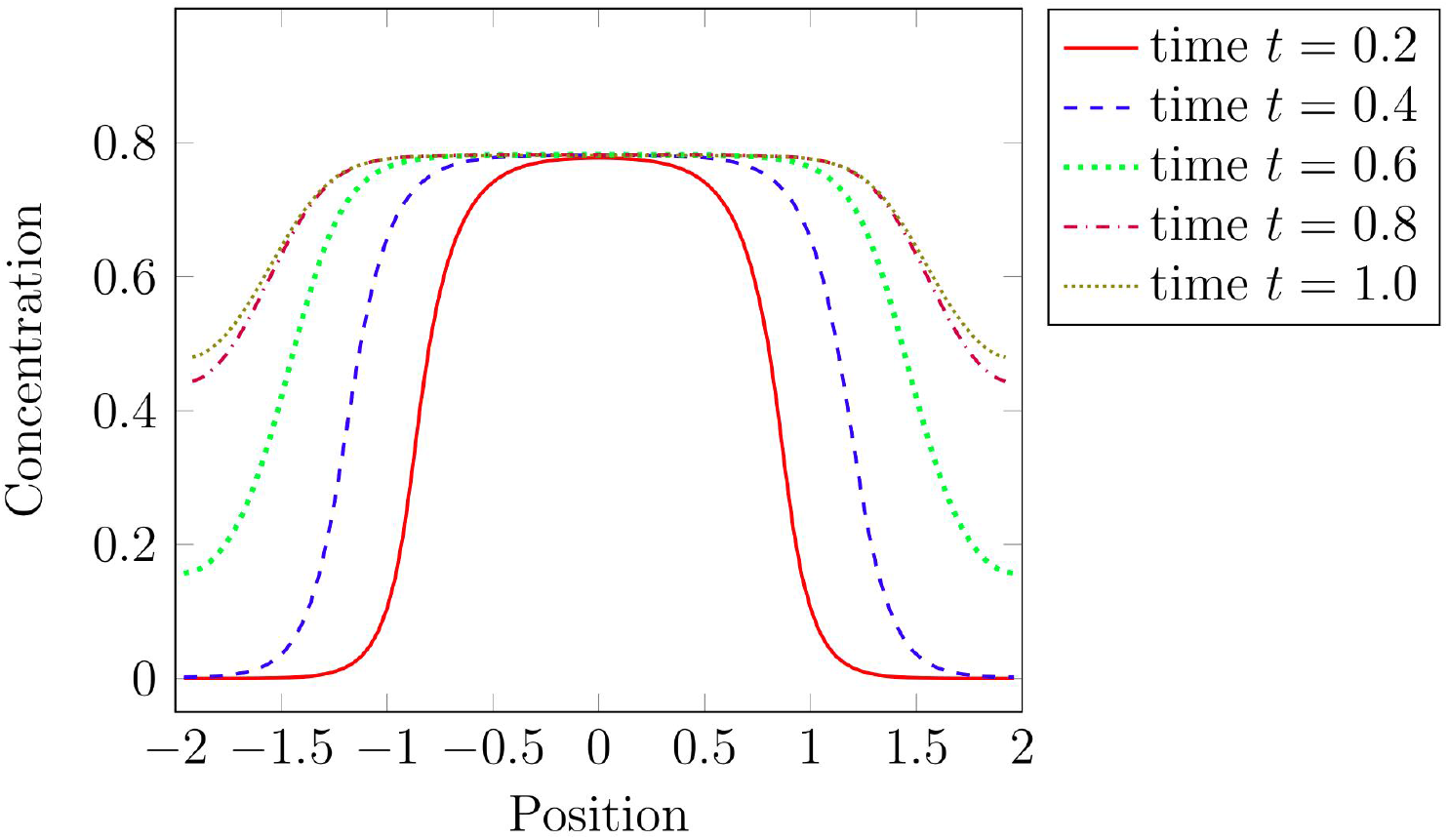
Time dynamics of CLV3 over-expression. Numerical calculation of CLV3 concentration for different time points after induction of ubiquitous WUS over-expression with *c* = 2.9: red solid for time *t* = 0.2, dashed blue for time *t* = 0.4, dotted green for time *t* = 0.6, dashdotted purple for time *t* = 0.8 and densely dotted olive time *t* = 1. As observed in experiments [42] the zone of high CLV3 expression extends in radial direction.

## Appendix A3. WUS loss of function

Fig. 11 shows time evolution of meristem and OC radius.

**Figure 11:**
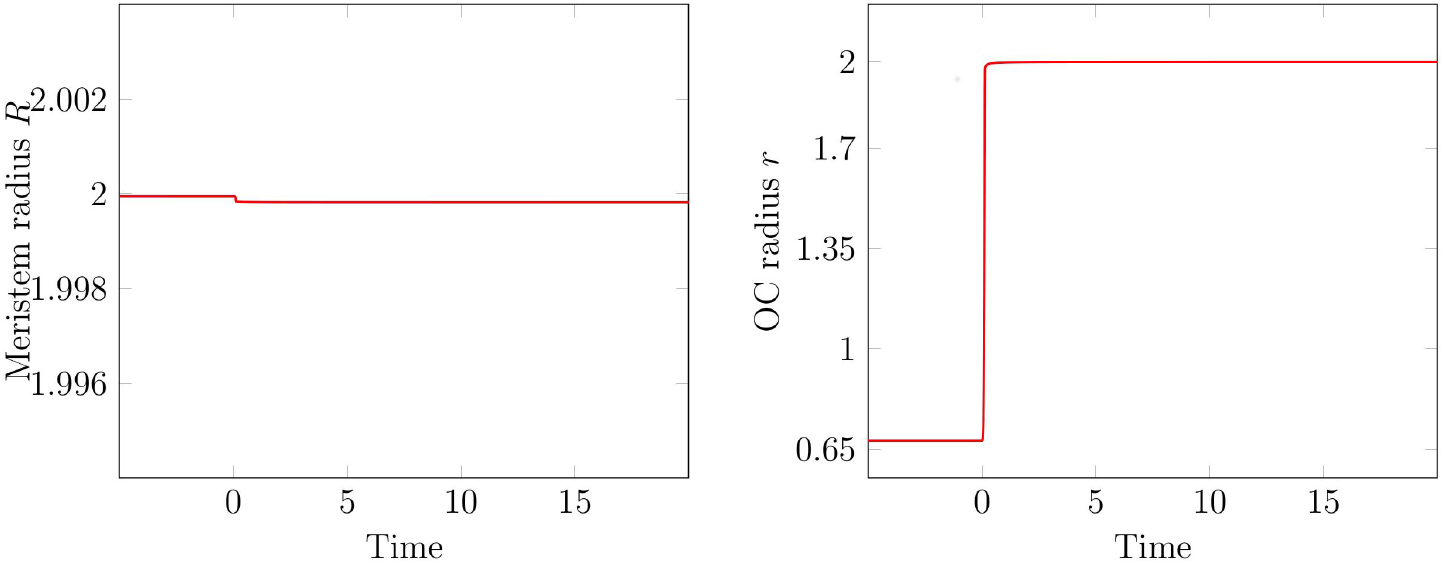
WUS loss of function. Time evolution of meristem and OC radius. WUS loss of function starts at time *t* = 0. The initial condition is the unperturbed equilibrium of the wild-type meristem.

## Appendix A4. CLV3 over-expression

Fig. 12 shows simulations for different levels of CLV3 over-expression. We observe that WUS concentrations are reduced compared to the wild type meristem. Different levels of over-expression lead to qualitatively similar results.

**Figure 12:**
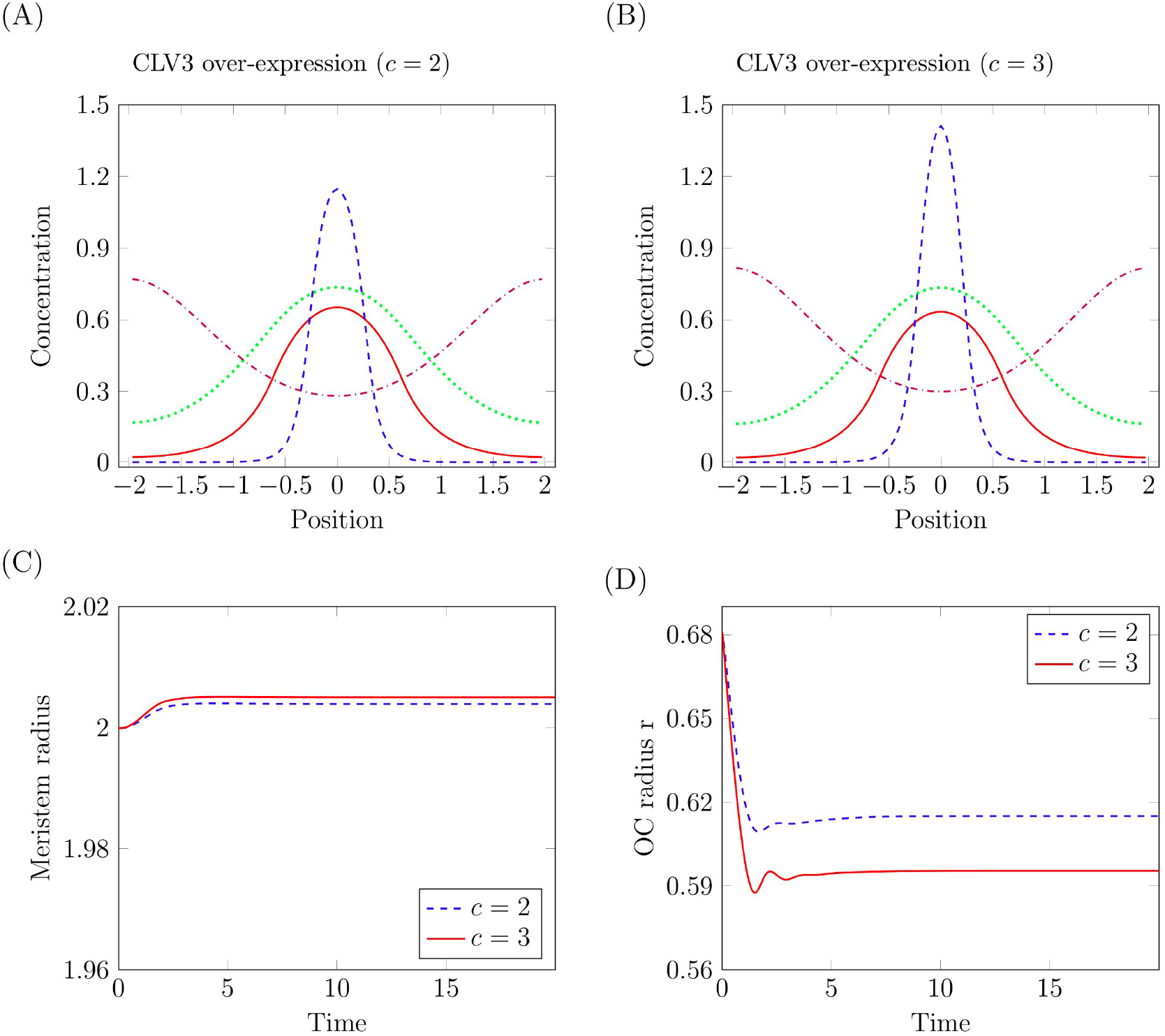
Different levels of CLV3 over-expression. Concentrations of WUS (red solid), CLV3 (dashed blue), CK (dotted green) and HEC (dashdotted purple) along the diameter of the meristem, position 0 corresponds to the center. CLV3 over-expression takes place in the cells which naturally express CLV3. It is simulated by multiplying the CLV3 production term by a constant *c*. (A) *c* = 2, (B) *c* = 3, (C-D) time evolution of meristem and OC radius after onset of over-expression.

## Appendix A5. CLV3 loss of function

Fig. 13 shows time evolution of meristem and OC radius. CLV3 loss of function leads to an increase of the OC radius.

**Figure 13:**
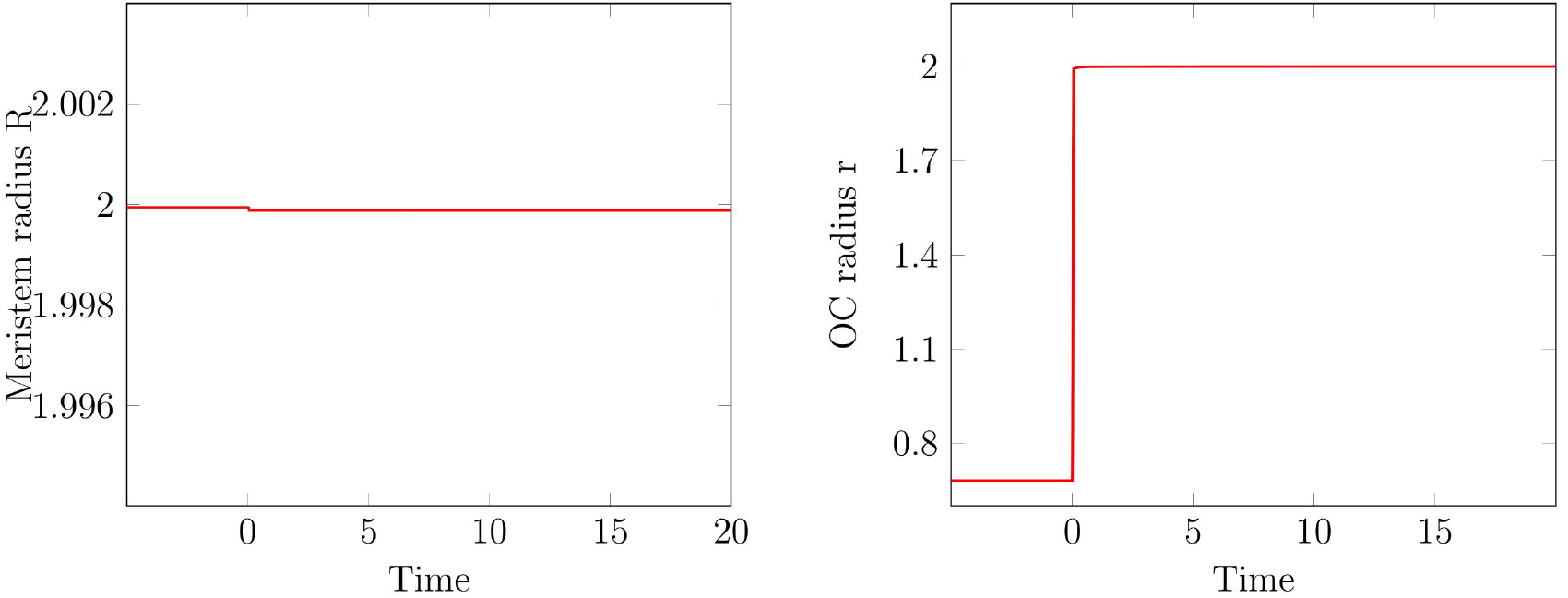
Meristem size in CLV3 loss-of-function. Time evolution of meristem and OC radius after CLV3 loss of function.

## Appendix A6. Reduced CK degradation

Fig. 14 shows simulations for different levels of reduction of CK degradation. Reduced CK degradation is modeled by the following modification of the equation for *u*_2_:

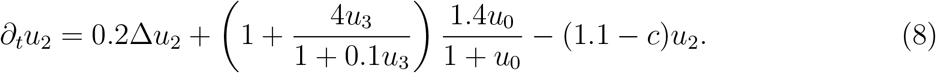

Reduced CK degradation leads to an increase of the OC radius. If the reduction is strong enough, we also observe an increase in meristem radius.

**Figure 14:**
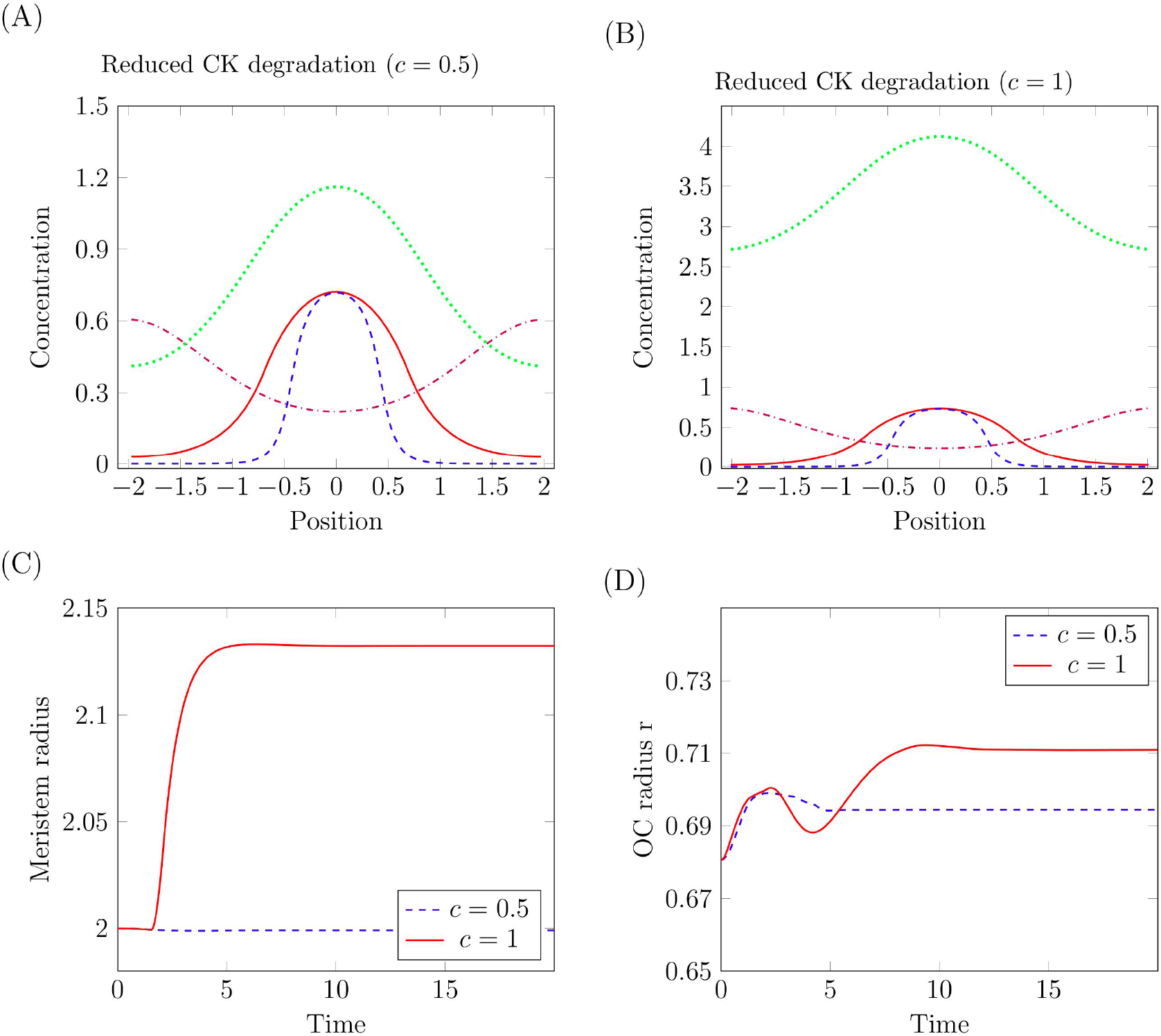
CK over-expression. Concentrations of WUS (solid red), CLV3 (dashed blue), CK (dotted green) and HEC (dashdotted purple) along the diameter of the meristem, position 0 corresponds to the center. CK over-expression takes place by changing the degradation rate of CK by a constant *c*. (A) *c* = 0.5, (B) *c* = 1, (C-D) time evolution of meristem and OC radius after onset of over-expression.

## Appendix A7. CK loss of function

Fig. 15 shows time evolution of meristem and OC radius in the case of CK loss of function. As observed in experiments meristem and OC radius decrease.

**Figure 15:**
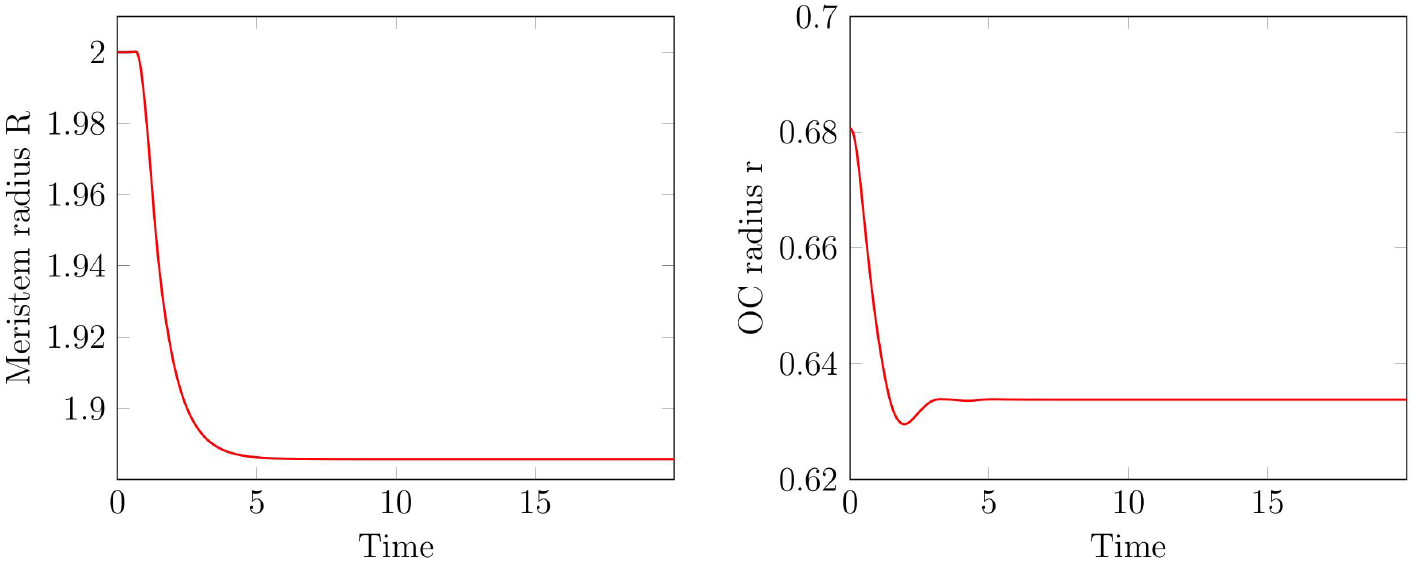
Meristem size in CK loss-of-function. Time evolution of meristem and OC radius in absence of functional CK.

## Appendix A8. HEC over-expression in stem cells

As in experiments HEC over-expression in stem cells leads to an increase of the meristem radius and the OC. The OC radius approaches the radius of the the total meristem, Fig. 16. Different levels of over-expression result in qualitatively similar dynamics. If we rerun the simulations omitting the direct effect of HEC on OC cell differentiation, the OC radius does not approach the meristem radius, see Fig. 7 (C-D) which is in contradiction to experimental observations [9].

To simulate latter scenario, in the ODE for *r* the HEC concentration was fixed to 0.3. This choice guaranties that the steady state for the modified model (without direct effect of HEC on OC cells) is identical to the steady state of the original model (with direct effect of HEC on OC cells).

**Figure 16:**
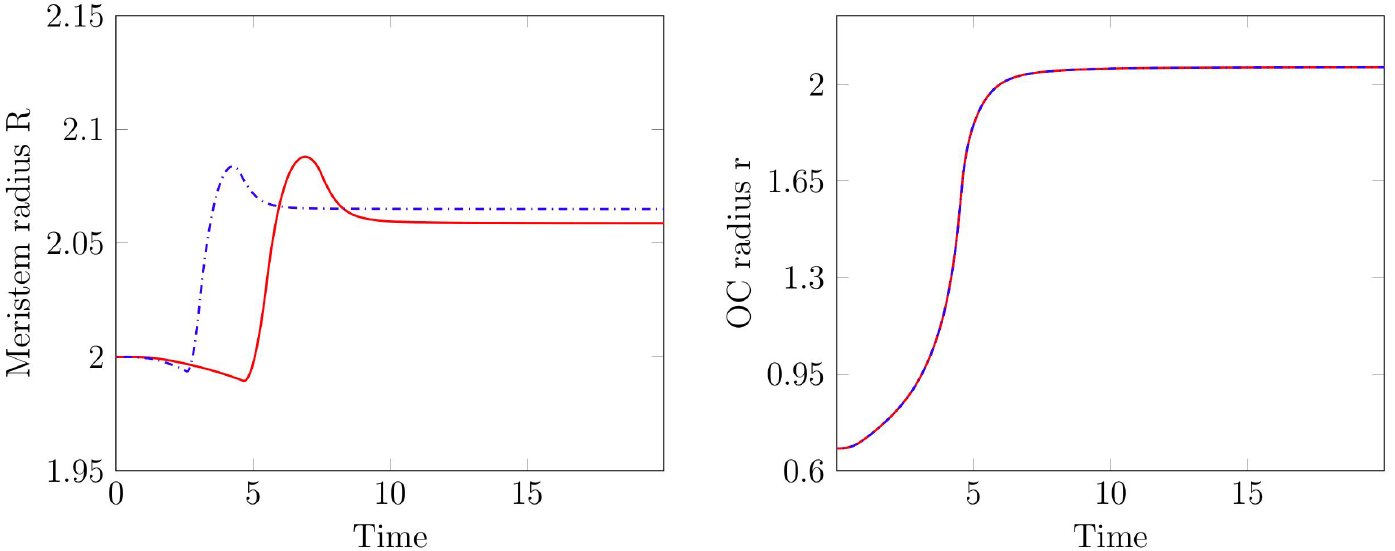
Meristem size in HEC over-expression. Time evolution of meristem and OC radius during HEC over-expression in stem cells. The level of over-expression is determined by the positive constant *c*. Results are shown for *c* = 0.05 (solid red) and *c* = 0.1 (dashdotted blue).

## Appendix A9. HEC loss of function

HEC loss of function leads to a smaller radius of the OC and the total meristem, see Fig. 17.

**Figure 17:**
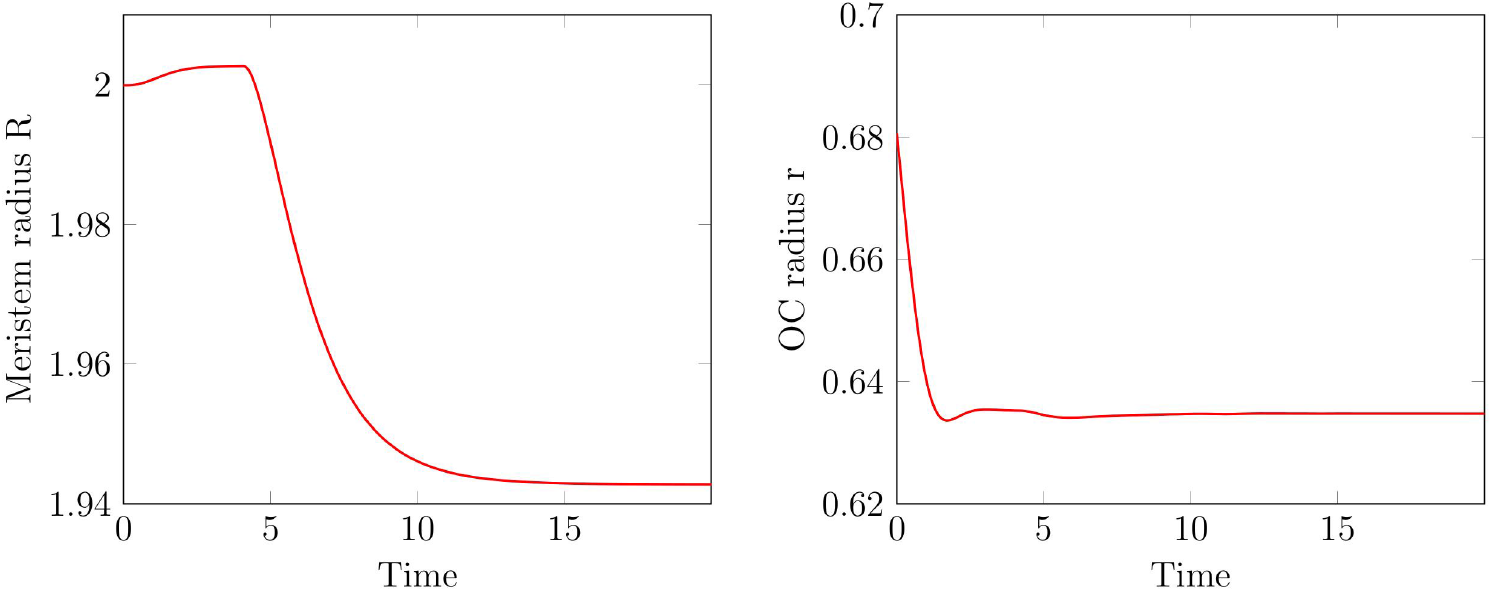
Meristem size in HEC loss of function. Time evolution of meristem and OC radius in absence of functional HEC.

## Appendix A10. Spatial dependence of proliferation rates

The term 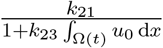 in equation (2) corresponds to the average proliferation rate in the SAM. To obtain the proliferation rate as a function of the radius we proceed as follows: We express the proliferation rate as a function of local WUS concentration, namely 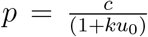. The parameter *k* is chosen such that the ratio between proliferation rate in the central and proliferation rate in the peripheral zone agrees with measurements from [35]. This implies *k* = 10. The constant *c* is chosen such that 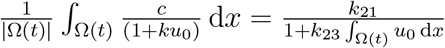. The obtained results are depicted in Figure 18.

## Cell differentiation

We calibrated our parameters so that the ratio of the radius of the CZ and the radius of the SAM is equal to the experimentally measured value. In the simulations, the CZ is defined by high levels of CLV3. We define all cells with a CLV3 level not less than 10% of the CLV3 level in the centre of the steady state meristem as stem cells. This definition is in line with the observation that approximately 10% of meristem cells are located in the CZ.

The model considers cell differentiation happening at the outer boundary of the SAM where cells leave the SAM and enter into organs.

The outflux rate of cells from the SAM into organs is given by the term 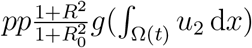 in equation (2).

**Figure 18:**
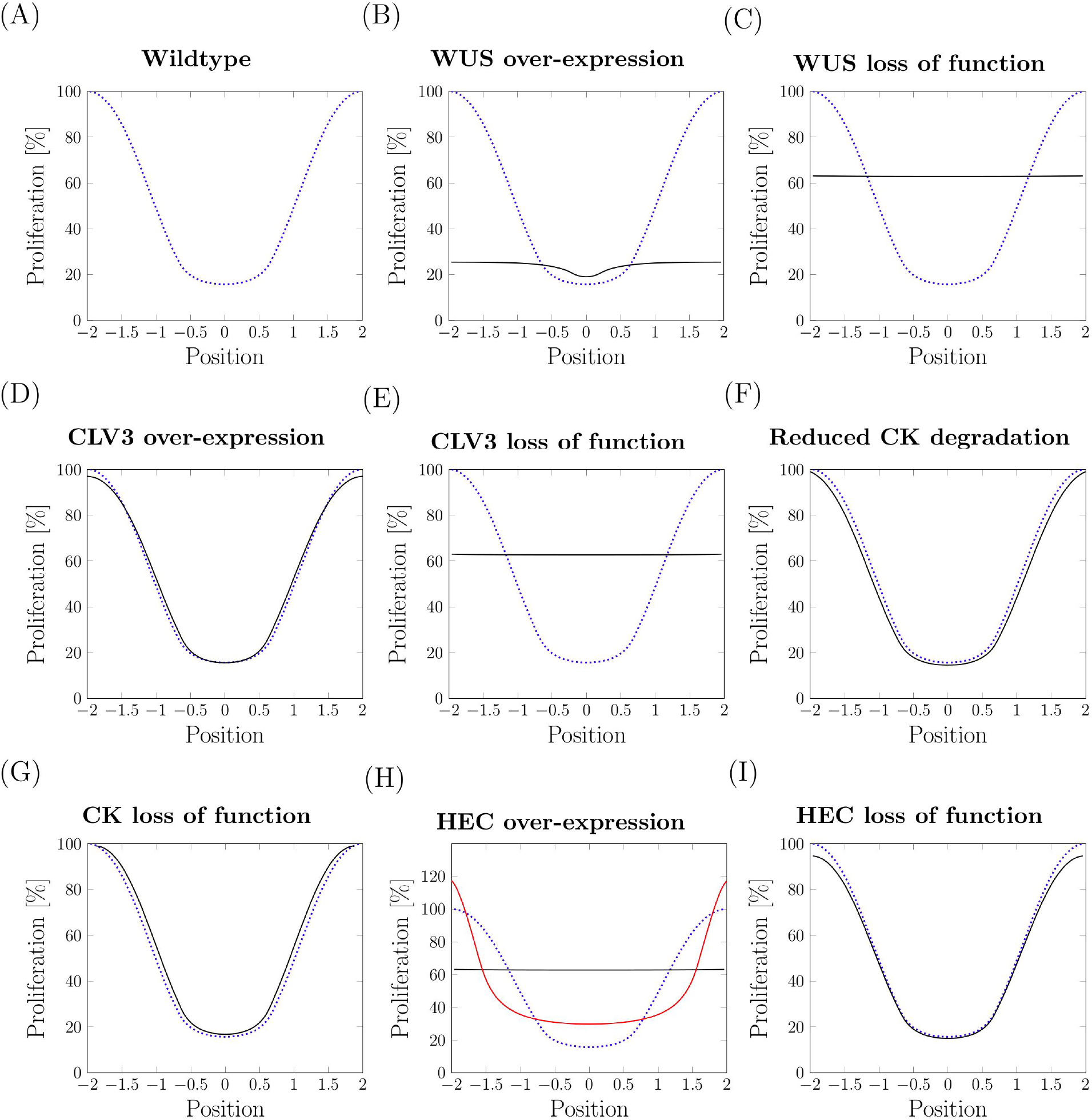
Spatial dependence of proliferation rates. Proliferation rate are shown as a function of the radial distance from the SAM center. They have been normalized with respect to the maximum rate in the wild-type steady state. The blue dotted line shows the proliferation rate of the wild-type equilibrium. The black lines show the proliferation rates at the end of the respective numerical experiments. The proliferation rates correspond system states shown in Fig 6. Dotted blue proliferation rate in steady state. (B) WUS over-expression *c* = 2 (C) WUS loss of function: red is with WUS=0 (D) CLV3 over-expression with *c* = 2 (E) CLV3 loss of function (F) Reduced CK degradation with *c* = 1 (G) CK loss of function (H) HEC over-expression with *c* = 0.05. The red line corresponds to the proliferation rate at the time when *r* = 0.75*R*. This is included because no measurements of proliferation rates at the end of the experiment are available for the HEC over-expression. (I) HEC loss of function.

## Appendix A11. Description of the numerical scheme

## Continuously deforming domains

We model the continuously deforming domain Ω(*t*) for time *t* > *t*_0_ as

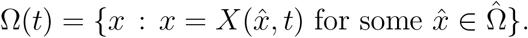

where 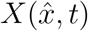 is the push-forward map from the reference domain 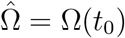 to time *t* > *t*_0_. The push-forward map is given by the parametric ordinary differential equation

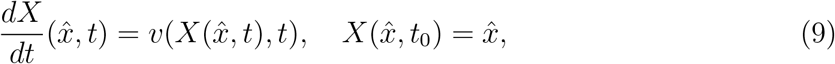

where *v*: Ω(*t*) × *t* → ℝ^*d*^ is a differentiable velocity field. Thus, the continuously deforming domain is given by the reference domain and the velocity field.

## Strong form of the reaction-diffusion equation

The reaction-diffusion equation results from conservation of mass. Let *ω*(*t*) ⊆ Ω(*t*) denote an arbitrary subdomain moving with the deformation (movement of points in *ω* is given by the push-forward map *X*). Conservation of a scalar quantity in *ω*(*t*) is stated by

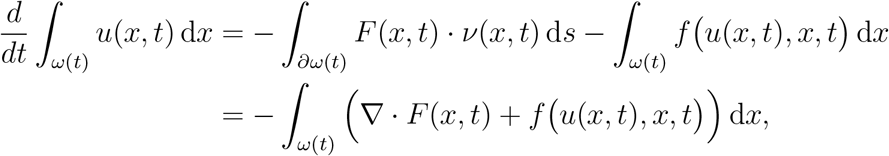

where *u*(*x, t*) is the density of the quantity to be conserved, *f* (*u, x, t*) is a source/sink/reaction term and *F* (*x, t*) is the flux of the conserved quantity given by Fick’s law

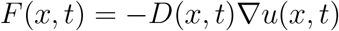

since *ω*(*t*) moves with the flow. Inserting the flux law and employing Reynolds’ Transport Theorem, we move the temporal derivative under the integral to obtain

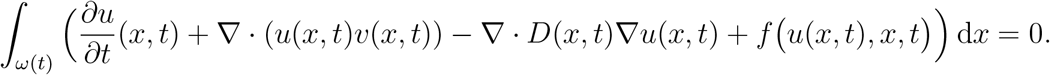

Since *ω*(*t*) was arbitrary the integrand must be zero pointwise (fundamental theorem of calculus) and we obtain the strong form of the reaction-diffusion equation on deforming domains:

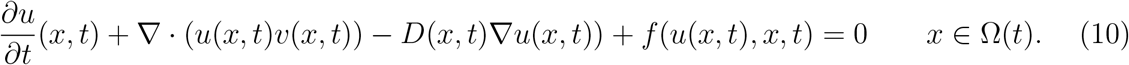

This equation is supplemented with initial conditions and boundary conditions which we assume to be of Neumann type for simplicity of notation:

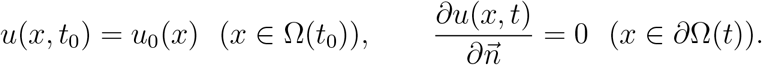

## Weak form of the reaction-diffusion equation

We first need to derive some auxiliary results. The inverse of the push-forward map is denoted by *X*^−1^(*x, t*). It maps point *x* at time *t* to a point in the reference domain. We assume the following identities hold (invertibility of push-forward map):

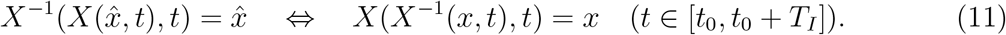

Differentiating the identity on the right by *t* and using the chain rule we obtain

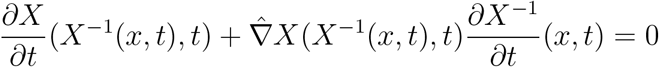

where 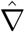 denotes the jacobian differentiating with respect to coordinates in the reference domain. Rearranging and using the definition of *X* from (9) we obtain

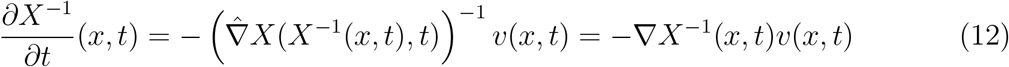

where used 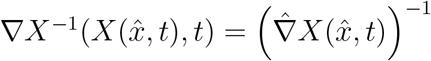 in the last step which is obtained from differentiating the left identity in (11) w.r.t. to all coordinates.

Next, consider a function *w*(*x, t*) on the continuously deforming domain by linking it to a function 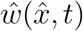 defined on the reference domain and any time, i.e.:

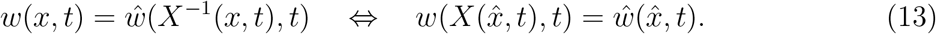

Applying the chain rule to the second identity we obtain the well-known transformation formula for gradients

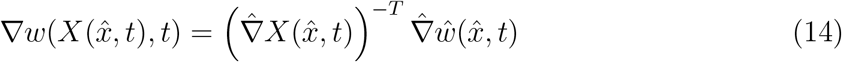

which allows to compute the gradient of a *w* on the deformed domain from the gradient of 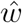 and the jacobian of the push-forward map at time *t*.

Applying the chain rule to the time derivative of the first identity in (13) and using the identities derived above yields the following important formula:

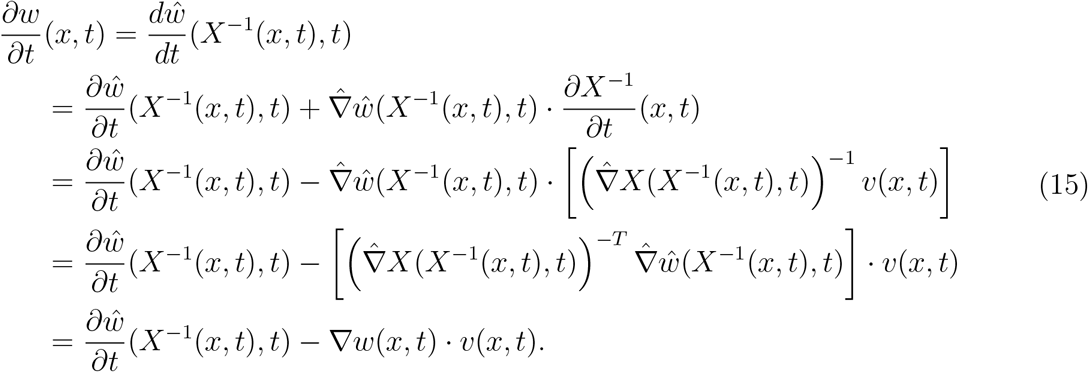

Using this result we derive a corollary of the Reynolds transport theorem. Formally applying the Reynolds transport theorem to the product *u*(*x, t*)*w*(*x, t*), where the dependence of *w* on time is assumed to be only due to domain deformation, i.e. 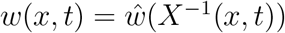, yields

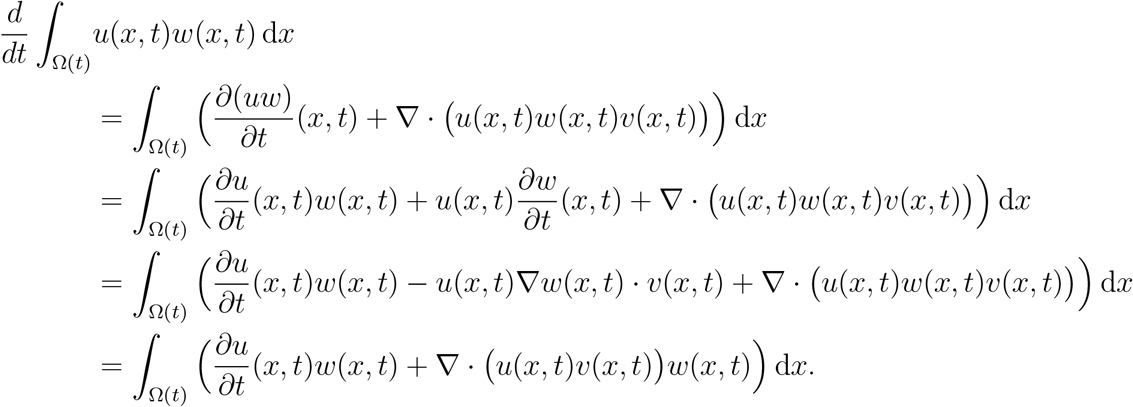

We are now in a position to derive the weak formulation of (10). Multiplying by a test function *w*(·, *t*) ∈ *H*^1^(Ω(*t*)) (depending on time only via the deformation of the domain), integrating by parts and using the corollary of Reynolds’ transport theorem yields:

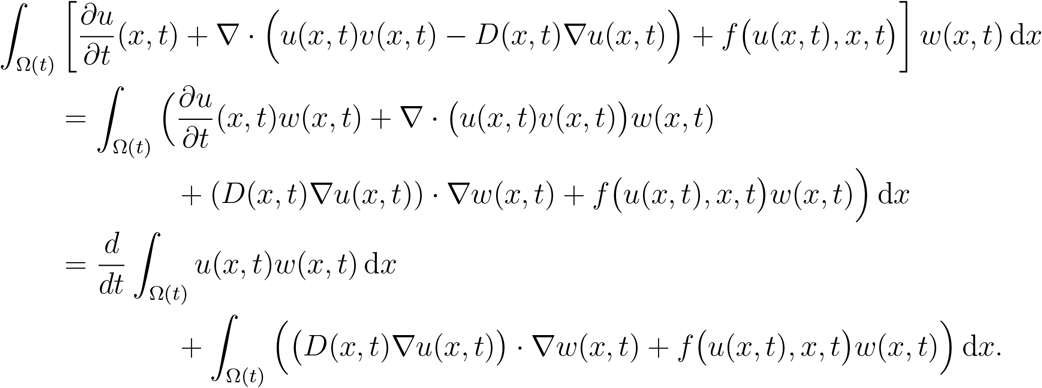

Setting *U* (*t*) = *H*^1^(Ω(*t*)) and *V* (*t*) = *H*^1^(Ω(*t*)), the weak formulation seeks *u* ∈ *L*_2_([*t*_0_, *t*_0_ + *T*]; *U* (*t*)) such that for all *t* ∈ (*t*_0_, *t*_0_ + *T*]

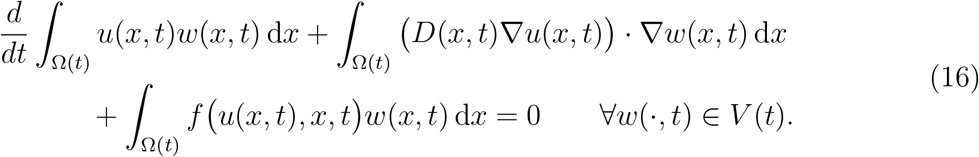

Observe that we chose to keep the temporal derivative outside of the integral.

## Finite element formulation

The FEM approximates the solution in a finite-dimensional subspace defined on a mesh. Here we assume that a mesh is obtained for the reference domain Ω(*t*_0_) which is then continuously deforming with the domain. The time-dependent mesh is denoted by its elements

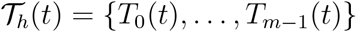

and the nodal positions

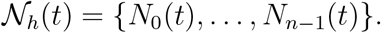

The connectivity of elements to nodes is given by the local to global map

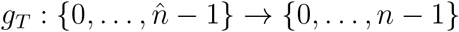

mapping the 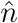 node numbers of the reference element 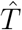 to global node numbers. The local to global numbering does not change throughout the computation and it is assumed that the quality of the mesh does not deteriorate with time (a reasonable assumption for the problem here).

Setting up the finite element problem is done by a pull-back to the reference element 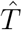 as usual. To that end define the map

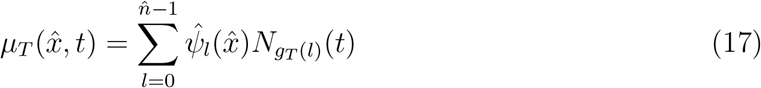

mapping a point 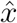 in the reference element 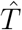 to the mesh element *T* at time *t*. Here, 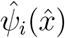 are the (multi-) linear Lagrange shape functions on the reference element. Consequently, the approximation Ω_*h*_(*t*) of the domain Ω(*t*) is polygonal (this could be easily extended to higher order elements but we refrain from doing so in order to keep the notation simple).

Now we are in a position to define the lowest order conforming finite element space on the continuously deforming mesh via the element transformation map:

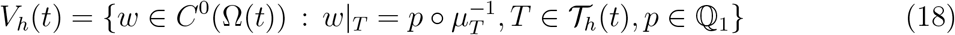

with 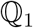 the multi-linear polynomials in dimension *d* (simplicial grids and piecewise linear can be used as well). For the test and ansatz functions we employ the subspace and affine subspace: *W_h_*(*t*) = *V_h_*(*t*) and *U_h_*(*t*) = *V_h_*(*t*). The time-continuous finite element problem now consists in finding *u*(*x, t*) ∈ *L*_2_([*t*_0_, *t*_0_ + *T*]; *U_h_*(*t*)) such that for all *t* ∈ (*t*_0_, *t*_0_ + *T*]

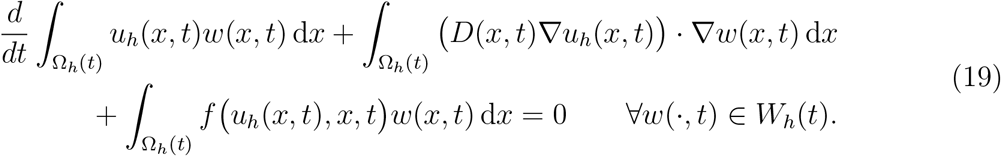

Choosing a basis representation

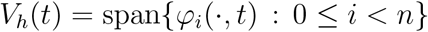

and inserting the ansatz

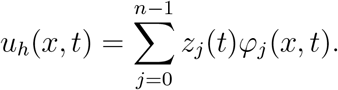

in (19) results in a system of ordinary differential equations (ODE) for the unknown coefficients *z_j_* (*t*).

For discretization in time introduce the possibly non-equidistant time steps

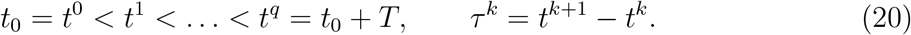

Denoting position of node *i* at time *t^k^* by 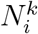 nodal positions are interpolated in time

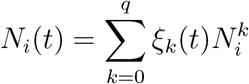

where the *ξ_k_* (*t*) are the one-dimensional, piecewise linear basis functions on the subdivision (20).

Given discrete nodal positions 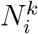 at *t^k^* and 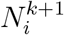 at *t*^*k*^+1, nodal positions are interpolated linearly

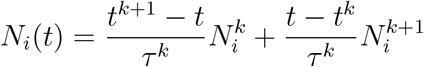

and we solve the system of ODEs using a second-order accurate diagonally implicit Runge-Kutta (DIRK) method. The required integrals are computed efficiently with the pull-back to the reference element given in (17).

## Operator splitting scheme for coupled problem

Above nodal positions 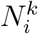 were given. We now treat the case where the nodal positions depend on the solution of the PDE. In the problem considered in this paper the domain is a circle with time-dependend radius *r*(*t*) centered at the origin. The change in radius is given a relation of the form

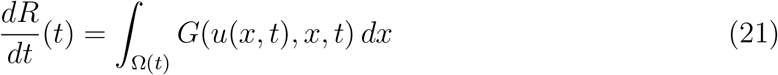

with some function *G*(*u, x, t*).

Coupling of the PDE and the domain growth is rather weak and we use first-order operator splitting scheme in time and explicit Euler for solving (21):

1. Generate mesh 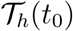 with nodal positions 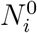 for Ω(*t*_0_) and initialize *u_h_*(*·, t*_0_) from initial value *u*_0_. Set *k* = 0 and choose a fixed time step *τ^k^* = *τ*.
2. Given nodal positions 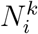 and *u_h_*(*·*, *t^k^*), compute a new radius via

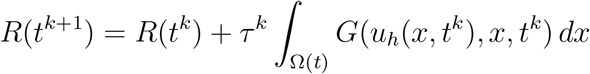

as well as new nodal positions

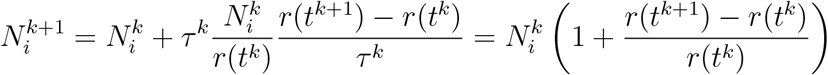

using the explicit Euler scheme.
3. Solve the PDE problem in (*t^k^*, *t*^*k*+1^] and go to 2.

## Appendix A12. Sensitivity

To study the sensitivity of the model we performed numerical experiments with perturbed parameters. In each simulation we perturbed all of the parameters by a random value from the interval [−*p, p*], where *p* was set to 2.5% and 5% of the respective parameter value. After 100 and 103 simulations (for *p* equal to 2.5% and 5%, respectively) we calculated the 95% confidence intervals (the 95% quantiles of the normal distribution with the mean value and standard deviation obtained from the sample). The obtained results are presented in Table 2. Obtained ratios between radii were normalized by division with the mean value of the sample. In Fig. 19 we present confidence intervals for *R*, *r* and the ratio between *r* and *R*. All quantities are divided by the mean values of the sample.

**Figure 19:**
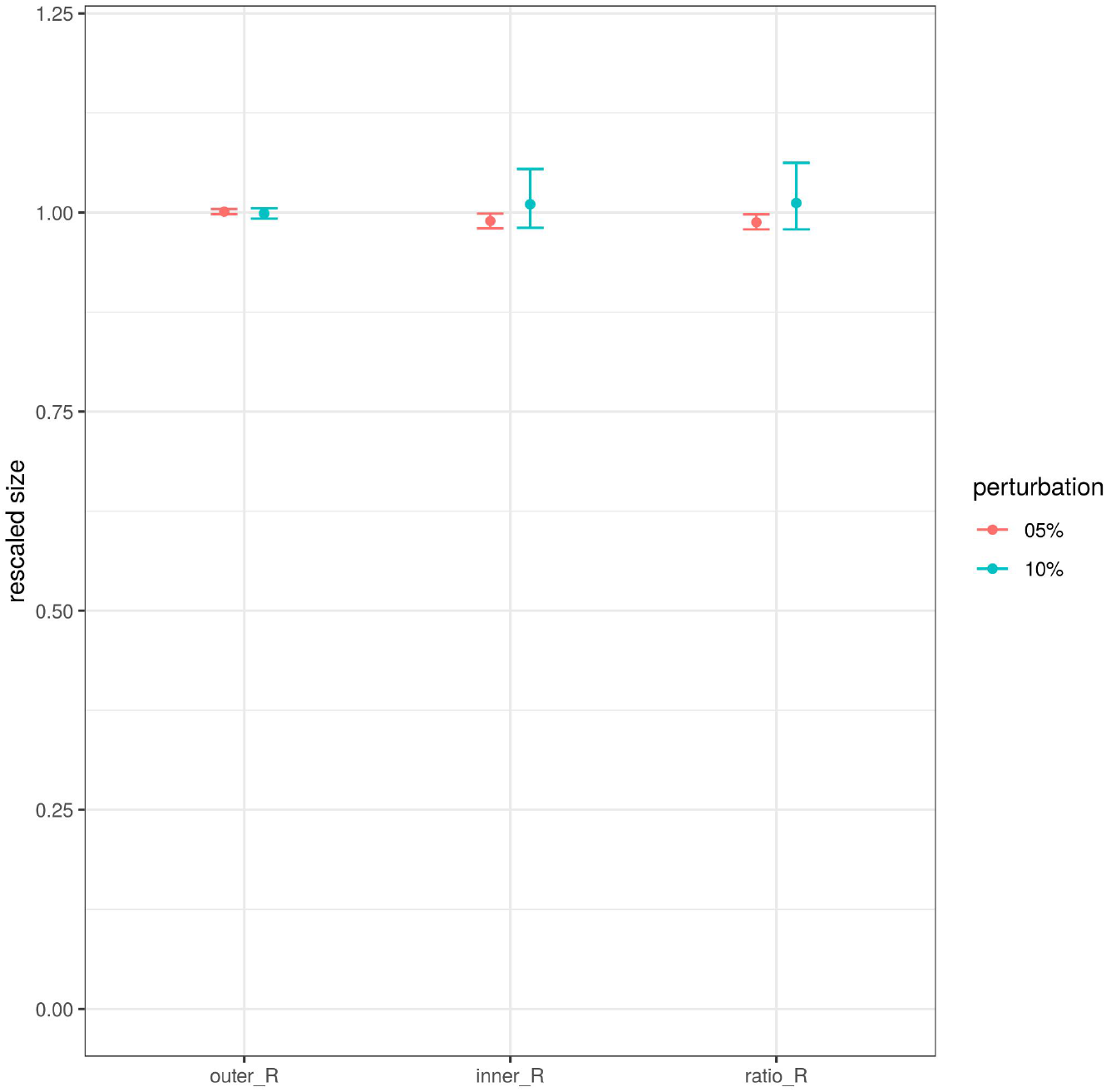
Sensitivity analysis. 95% confidence interval for perturbation ±2.5% (red line 05%) and ±5% (blue line 10%). All values are presented on rescaled axes.

All simulations were run on the mesh *N* = 50 (90404 degrees of freedom) with time step *dt* = 0.05. The variation in the ratio *r/R* is of the same order of magnitude as the perturbation of the parameters. This implies that the model is robust with respect to perturbations of parameter values.

## Acknowledgments

This work was supported by research funding from the German Research Foundation DFG (Collaborative Research Center SFB 873, Maintenance and Differentiation of Stem Cells in Development and Disease).

